# The actin cortex acts as a mechanical memory of morphology in confined migrating cells

**DOI:** 10.1101/2024.08.05.606589

**Authors:** Yohalie Kalukula, Marine Luciano, Guillaume Charras, David B. Brückner, Sylvain Gabriele

## Abstract

Cell migration in narrow microenvironments is a hallmark of numerous physiological processes, involving successive cycles of confinement and release that drive significant morphological changes. However, it remains unclear whether migrating cells can retain a memory of their past morphological states, which could potentially enhance their navigation through confined spaces. By combining cell migration assays on standardized microsystems with biophysical modeling and biochemical perturbations, we demonstrate that local geometry governs these morphological switches, thereby facilitating cell passage through long and narrow gaps. We uncovered a long-term memory of past confinement events in migrating cells, with morphological states correlated across transitions through actin cortex remodeling. These findings suggest that mechanical memory in migrating cells plays an active role in their migratory potential in confined environments.

---

Adherent cells actively adapt their shape and functions to the physicochemical constraints imposed by their surrounding extracellular matrix (ECM) in response to varying ECM properties(*1–3*). A crucial facet of this adaptive response is the concept of mechanical memory, wherein a new phenotype persists even when the physical constrains are relaxed. Emerging evidence suggests that adherent cells can retain a memory of the mechanical characteristics of their microenvironment after prolonged exposure, ranging from a few days to several weeks, through changes in transcription factor activity and epigenetic modifications(*4–7*). Despite these observations, the concept of mechanical memory related to short duration confinement events that occur repetitively and frequently over hour-long periods during confined migration remains unexplored. In physiologically relevant contexts, cells must interact dynamically with their surroundings to migrate and facilitate processes such as embryogenesis(*8*), immune response(*9*), wound healing(*10*), and cancer metastasis(*11*). Migrating cells must therefore adapt their strategies to navigate through mechanically heterogeneous matrices and narrow constrictions, leading to variable confining and unconfining events associated with significant morphological switches. Since adaptation to confinement is potentially energetically costly due to the necessity for transcription and translation, retaining a long-term memory of past confinement may provide a significant advantage for migrating cells.

## A morphological switch facilitates passage through long narrow gaps

To study cell shape changes during confined migration under standardized and reproducible conditions, we developed a system of adhesive micropatterns. This system features a dumbbell geometry(*12*) with a narrow fibronectin (FN) passage of W=6 microns in width, with variable lengths ranging from L=40 to 160 microns **(Fig. 1A)**. These dimensions correspond to those of interstitial spaces found for instance in skin(*13*) and breast tissues(*14*), which are less than 10 µm in diameter and can extend over 150 µm in length(*15*). Each end of the narrow bridge is connected to a large square area (40x40 µm), allowing the cell to spread and repolarize before traversing the confined passage again **(Movie S1).** Interestingly, it has been demonstrated that one-dimensional (1D) microstripes, similar to those used for the bridge, can effectively replicate many characteristics of the rapid, uniaxial migration phenotype observed in fibrillar three-dimensional (3D) cell-derived matrices(*16*). Quantifying the cell shape index (CSI, see methods) (*17*) of mammary epithelial cells (MCF-10A) within these confined bridges **(Fig. 1B)** based on automatic tracking of 20 hours time-lapse experiments (**Movie S2)** revealed a surprising bimodal distribution for the longest length (160 microns, **Fig. 1C)**. This suggests the presence of two distinct sub-populations of cell morphologies with CSI either below or above 0.4 (**Fig. 1D and Movie S3)**, corresponding to elongated (≈38%) or compacted (≈62%) morphologies, respectively **(Fig. 1E)**. Furthermore, not all crossings were successful, and success appeared correlated with CSI. To quantify this observation, we tracked back-and-forth motions by time-lapse imaging for the different bridge lengths. This enabled quantification of the percentage of failed passages **(Fig. 1F and Movie S4)**, during which cells reversed direction before reaching the square at the opposite end, versus successful passages **(Fig. 1G and Movie S4)**, during which cells successfully traversed the entire bridge before spreading in the opposite square. Surprisingly, we observed that cells crossed longer bridges more efficiently: the probability of an attempt (defined as a protrusion entering the bridge) to result in a successful transition increased with bridge length, reaching ≈58.2% on the longest bridges of 160 µm. Surprisingly, this high rate of successful crossings is directly linked to the compacted morphology **(Fig. 1H)**, which appears to be much more effective for navigating through long and narrow environments **(Fig. 1I)**.

**Figure 1.**
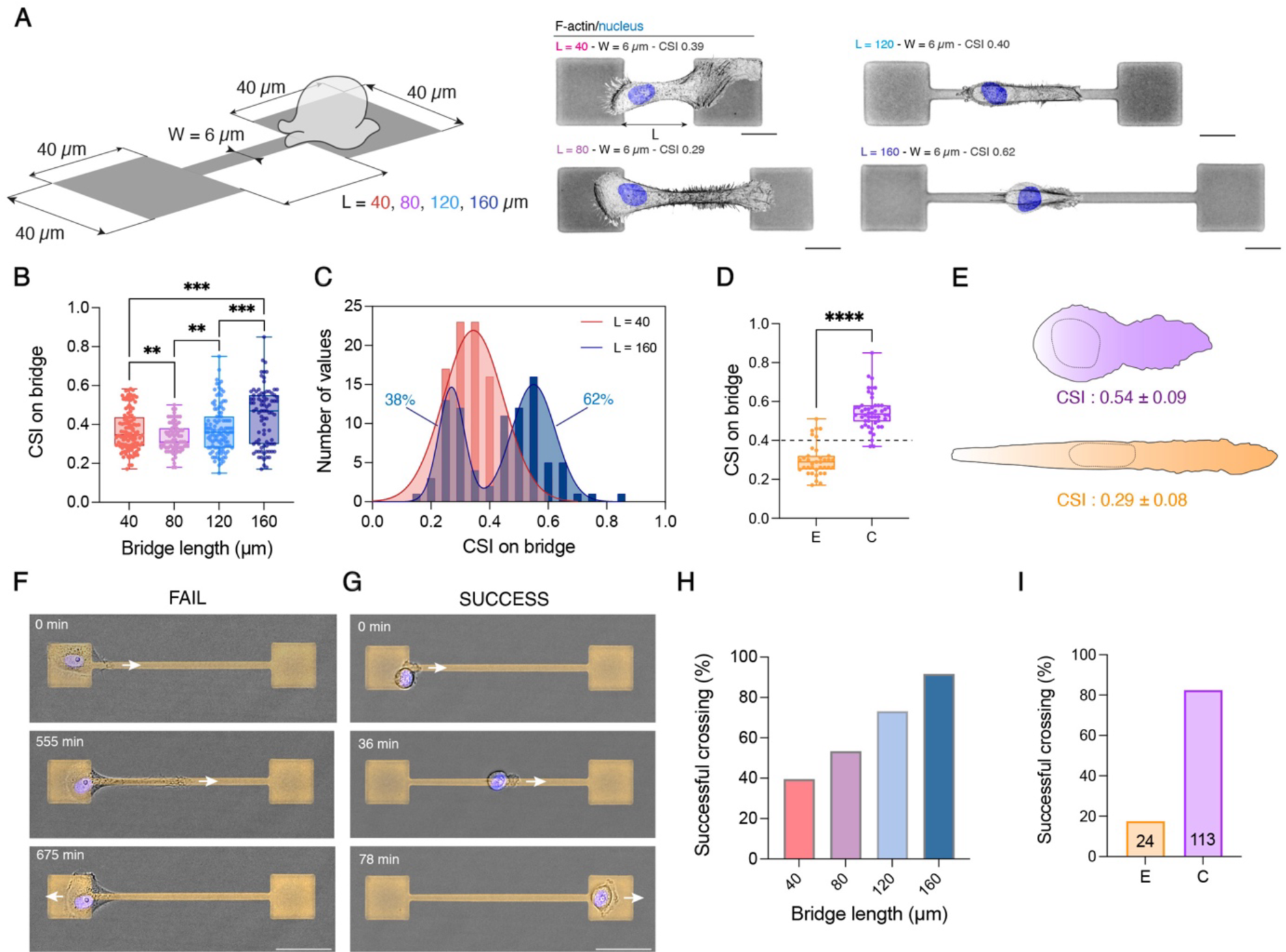
A morphological switch ensures a successful crossing on long bridges. **(A)** Schematic representation and typical microscopy images of single epithelial cells (MCF-10A) navigating through a narrow passage represented by a dumbbell micropattern. The fibronectin-coated micropattern consists of two square islands of 40 μm × 40 μm connected to a narrow bridge of a constant width (W = 6 µm) and varying lengths (L= 40, 80, 120 and 160 µm). The nucleus is stained in blue with Hoechst and actin filaments are stained with phalloidin with the color inverted. The cell shape index (CSI) is indicated for each bridge length. Scale bars, 20 µm. **(B)** Cell shape index on bridge versus bridge length (n = 116 for 40 µm, n=76 for 80 µm, n=109 for 120 µm and n=87 for 160 µm from N≥3). **(C)** Distribution of the Cell shape index on bridge for L=40 µm (in red) and L=160 µm (in blue). **(D)** Cell shape index on bridge for elongated (n=37, orange) and compacted (n=50, purple) epithelial cell morphologies on L=160 µm. **(E)** Schematic representation of the typical morphology of a compacted (CSI: 0.54±0.09, purple) and an elongated (CSI: 0.29±0.08, in orange) morphology. Time-lapse sequence of **(F)** a failed and **(G)** a successful crossing on a bridge of W=6 µm and L=160 µm. White arrows show the direction of migration. Scale bar, 50 µm. Percentage of successful crossings for **(H)** bridges of varying lengths (n = 116 for 40 µm, n=76 for 80 µm, n=109 for 120 µm and n=87 for 160 µm from N≥3 replicates per condition) and **(I)** elongated (n=24) versus compacted (n=113) morphologies on L=160 µm. **p < 0.01, ***p < 0.001, ****p < 0.0001.

## Symmetry breaking and polarization state drive the dynamics of elongated and compacted morphologies

To explore the link between cell morphology and confined migration efficiency— measured by the percentage of successful crossings—we quantified back-and-forth cell movements over a 20-hour period and color-coded the trajectories based on CSI **(Fig. 1D-E)**, revealing striking differences in migration dynamics between the two morphologies **(Fig. 2A)**. Compacted cells exhibited an average crossing speed three times faster than elongated morphologies **(Fig. 2B)** and three times shorter dwell times—defined as the time spent on the square areas—compared to elongated cells (**Fig. 2C)**. Actin flow analysis in the lamellipodia **(Fig. 2D and Movie S5)** of both morphologies revealed a significantly higher retrograde flow in compacted cells **(Fig. 2E-F)**, a hallmark of fast-migrating cells (*18*). Interestingly, morphological switches from an elongated to a more compacted state have been previously observed during the migration of breast epithelial cells in collagen microtracks with a high level of spatial confinement(*19*). Furthermore, the intrinsic relationship between adopting a compacted mode and the increase in migration speed has been observed for other cell types in different confinement situations, such as migration on glass fibers with a diameter similar to the bridge(*20*) width **(Supp. Fig. S1A)** or within collagen matrices(*21*) **(Supp. Fig. S1B)**. High-resolution confocal microscopy **(Fig. 2G and Supp. Movies S6-S7)** revealed that elongated cells appeared symmetric, with no distinct difference between the front and rear **(Fig. 2G, top)**. In contrast, compacted cells exhibited the characteristic morphology of migrating cells, with a spread leading edge and a rounded rear **(Fig. 2G, bottom)**. Compacted cells were characterized by significantly smaller spreading areas **(Fig. 2H)** and a lower total amount of actin filaments than their elongated counterparts **(Fig. 2I)**. Intriguingly, compacted cells displayed marked asymmetry in actin filament arrangement **(Fig. 2J)**, demonstrating a pronounced break in symmetry characteristic of the strong cell polarization state in fast migratory cells(*22*). To test the hypothesis that the difference in migratory dynamics between compacted and elongated cells may originate from a symmetry-breaking event in cell polarity, we explored a minimal biophysical model of active migration under confinement. Specifically, we modelled cells as polar particles exerting active migration forces in a direction of polarity ***p*** = (*p_x_*, *p_y_*). The dynamics of cell position *x*(*t*) are then described by d*x*/d*t* = *p_x_* + *F*(*x*), where *F*(*x*) are forces arising due to the confinement by the micropattern boundaries **(Supp. Theory Note)**. To describe the dynamics of cell polarity, we used a minimal model that allows both states in which cells are unpolarized (corresponding to morphologies with two opposing protrusions), and states in which cells are highly polarized in the direction of a single protrusion. Specifically, we hypothesized that the dynamics of cell polarity follow ***p*** = (−*β*|***p***|^2^ − *α*)***p*** + *γ**F***(*x*) + *σξ*(*t*) with *β* > 0, where *γ* quantifies the tendency of cells to repolarize upon contact with the micropattern boundary, and ξ(*t*) is a Gaussian white noise modeling fluctuations of the cell polarization. Different states of polarization are determined by the parameter *α*: for positive *α*, the cell polarization stochastically fluctuates around a mean-zero polarization state **(Fig. 2K, top)**. For negative *α*, the cell is highly polarized, with a non-zero mean polarity **(Fig. 2K, bottom)**. Simulations of our model with positive and negative *α* produced trajectories **(Movie S8)** in close agreement with those observed for elongated and compacted cell morphologies, respectively **(Fig. 2L and Movie S3)**. Fitting *α*-values to the experimentally measured crossing speeds, we indeed find positive and negative *α* for elongated and compacted cells, respectively **(Supp. Theory Note)**. Our model correctly predicts that compacted cells were faster **(Fig. 2M)** and spend less time on the square islands between transitions, as quantified by the dwell time **(Fig. 2N)**. Furthermore, our model makes a key prediction for the qualitative nature of the nonlinear dynamics of elongated and compacted cells during their transition across the bridge. Elongated cells should exhibit a stable fixed point at vanishing speed *v* = 0, indicating that they are susceptible to stochastic fluctuations, leading to frequent changes in direction **(Fig. 2O)**. Compacted cells exhibit two stable fixed points a finite speed, leading to a highly persistent crossing of the bridge at a fixed speed. These predicted dynamics can be measured experimentally by inferring the cell’s average acceleration as a function of its velocity of elongated and compacted cells(*23*) **(Fig. 2P, Supp. Theory Note)**. This result confirmed the hypothesized qualitative difference in nonlinear dynamics, with compacted cells exhibiting a pronounced separation of two stable fixed points at finite speeds *v* ≈ 2 *μ*m/min, close to the measured typical crossing speed. Together, our findings demonstrate that these morphological states control distinct migratory dynamics due to distinct polarization states of confined cells.

**Figure 2.**
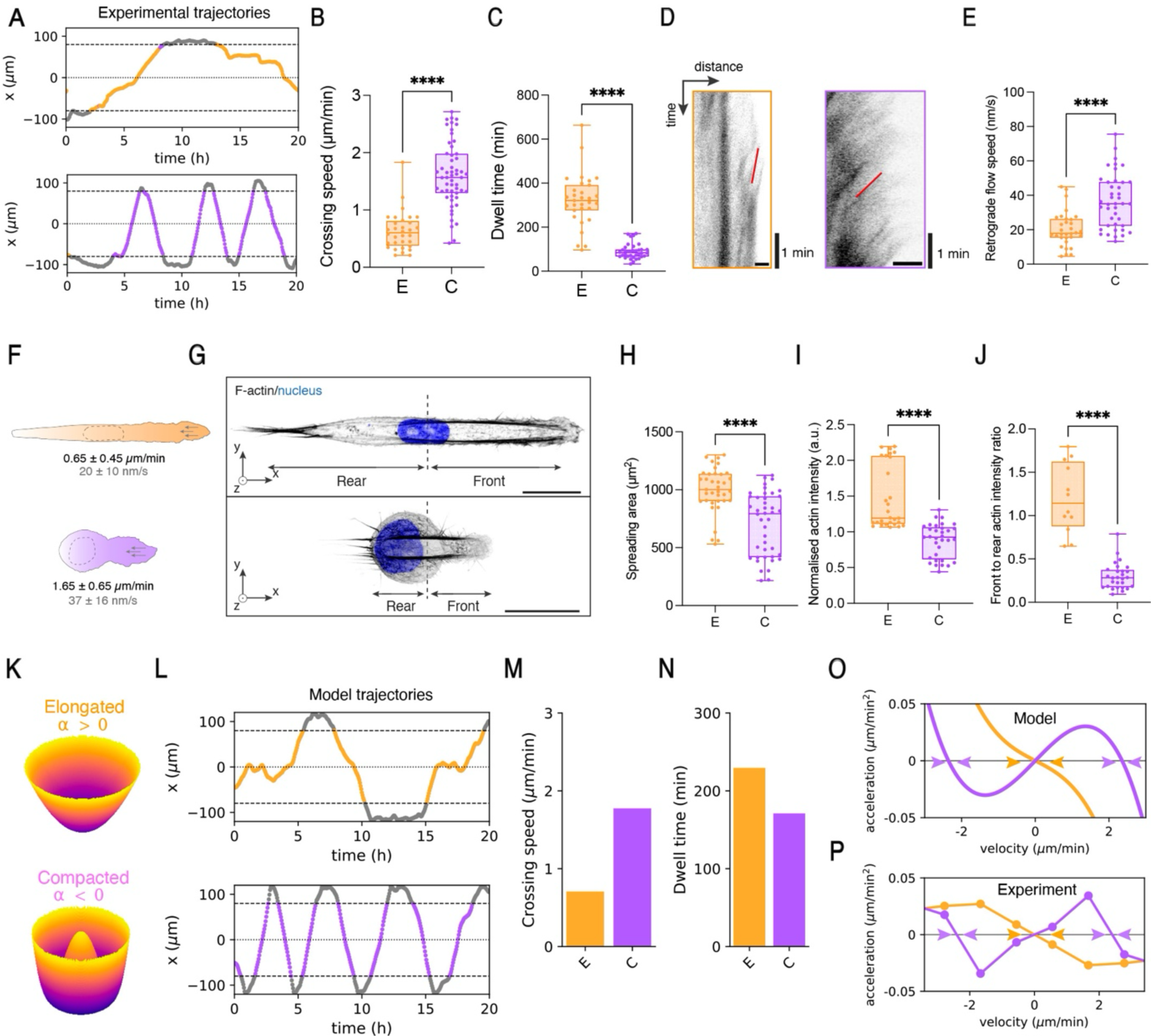
Impact of symmetry breaking and polarization states on cell dynamics. **(A)** Representative experimental trajectories for an elongated (top) and a compacted (bottom) cell morphology on a micropatterned dumbbell of 160 µm long and 6 µm wide. Trajectories on the bridge are color-coded with elongated morphologies in orange (CSI<0.4) and compacted morphologies in purple (CSI>0.4), while square zones are in grey. **(B)** Crossing speed (n=32 for elongated, n=55 for compacted, and N=16 replicates per condition) and **(C)** dwell time for elongated (n=31 and N=16, orange) and compacted (n=36 and N=16, purple) cell morphologies. **(D)** Kymographs of the actin flow and **(E)** quantification of the retrograde actin flow speed in the lamellipodia elongated (in orange) and compacted (in purple) cells. **(F)** Representative sketch of the elongated (n=32 and N=2, orange) and the compacted (n=41 and N=2, purple) morphologies with their associated mean cell speed (in black) and retrograde actin flow speed (in grey). Grey arrows show the direction of the retrograde flow. **(H)** Spreading area (n=37 for elongated, n=42 for compacted, N=2). **(I)** normalized actin density (n=28 for elongated, n=33 for compacted, N=2) and front-to-rear actin intensity ratio (n=12 for elongated, n=28 for compacted, N=2) for elongated (in orange) and compacted (in purple) cell morphologies. **(K)** Representation of the cell’s polarity dynamics determined by the parameter, *α*, with *α* > 0 when the cell polarization stochastically fluctuates around a mean-zero polarization state (top), and *α* < 0 when the cell is highly polarized (bottom), with a non-zero mean polarity. **(L)** Representative trajectories, **(M)** crossing speed and **(L)** dwell time predicted by the model with elongated morphologies in orange, compacted morphologies in purple and the time spent on squares in grey. Average acceleration of elongated (in orange) vs compacted (in purple) cells as a function of the cell velocity as **(O)** predicted by the model and **(P)** measured experimentally (n=57 cells and N=16 replicates per condition). Purple arrows show the presence of stable fixed points at finite speeds *v* ≈ 2 *μ*m/min close to experimental crossing speed values, while orange arrows show *v* = 0 where both curves cross each other. ****p < 0.0001 and n.s. not significant.

## Dynamic morphological switch is controlled by local geometry

Importantly, single cells migrating in a confined environment can alternate between elongated and compacted morphologies. This leads to morphological switch events during migration **(Fig. 3A, Movie S9)**, which are accompanied by an immediate change in migration speed, consistent with our previous results **(Fig. 2B).** This raises a central question: what are the statistical rules of cells switching between these two distinct morphological states? To gain insight into this, we analyzed the evolution of cell shape over time during crossing events **(Fig. 3B)**, using numerous experimental trajectories extracted from 20-hour time-lapse movies. Based on the CSI, we automatically assigned in these trajectories the time spent in the elongated mode (orange), compacted mode (purple), and on the deconfinement squares (gray) **(Fig. 3C)**. Interestingly, we found that the percentage of compacted cells, defined as having an average CSI on the bridge greater than 0.4 **(Fig. 1D)**, strongly increased with the bridge length, reaching ≈58% for a bridge length of 160 um **(Fig. 3D)**.

**Figure 3.**
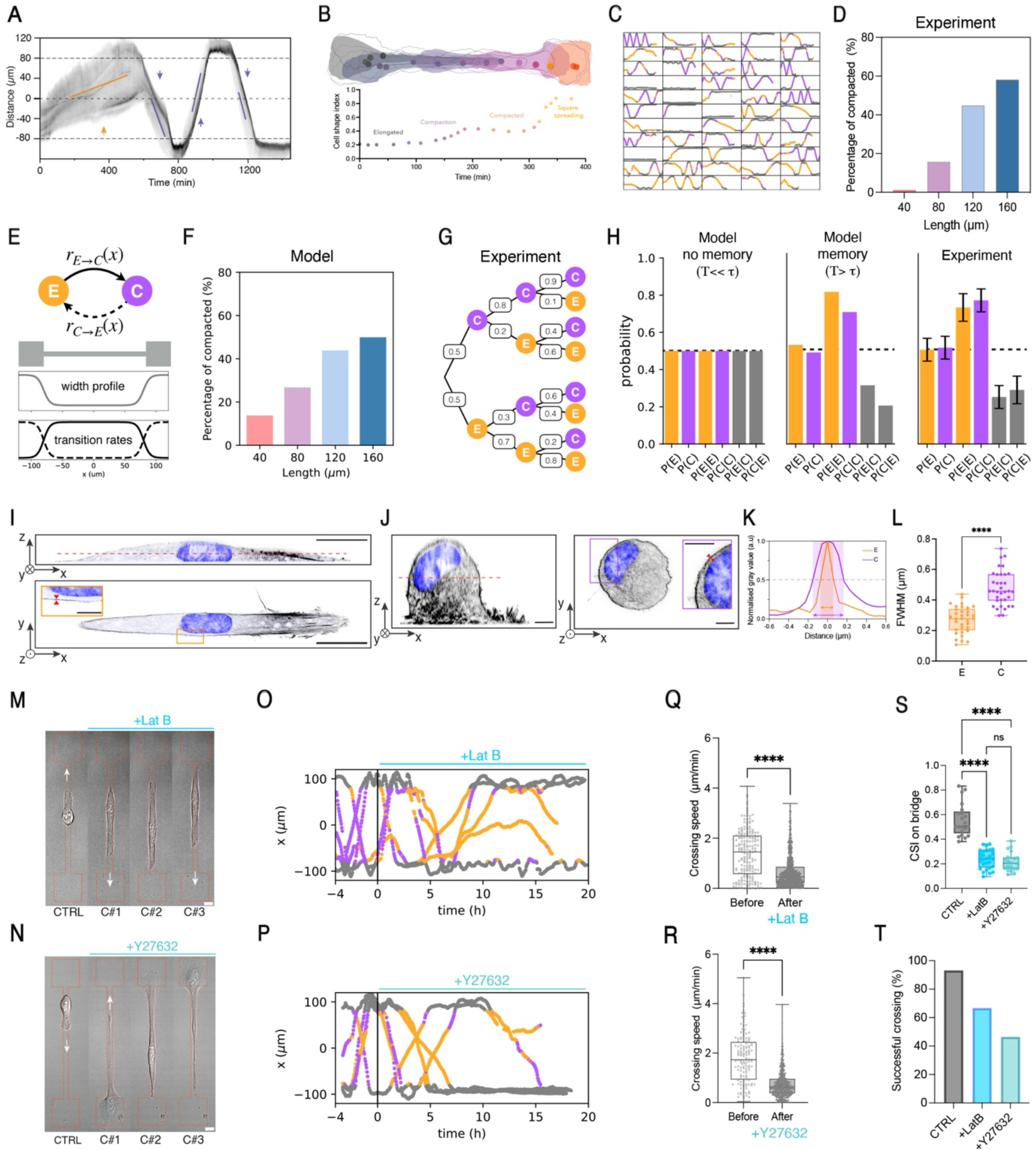
Morphological switch dynamics and mechanical memory. **(A)** Representative kymograph of a migrating cell tagged for actin with Spy555-FastAct, showing a transition from an elongated to a compacted state over a 24-hour time lapse. **(B)** Representative color-coded outline of an MCF-10A cell undergoing a morphological switch on a dumbbell micropattern, alongside the corresponding evolution of the CSI vs time. **(C)** Selection of n=55 color-coded cell trajectories of individual MCF-10A cells migrating on FN dumbbell micropatterns for 20 hours. Trajectories on the bridge are color-coded: elongated morphologies in orange (CSI<0.4) and compacted morphologies in purple (CSI>0.4), with square zones in grey. **(D)** Percentage of compacted cells for varying bridge lengths (n = 116 for L=40 µm, n=76 for 80 µm, n=109 for 120 µm and n=87 for 160 µm from N≥3 replicates per condition). **(E)** Schematic representation of the geometry-sensitive stochastic switching between elongated and compacted cells, including the spatial dependence of the normalized switching rates. **(F)** Theoretical prediction of the percentage of compacted cells for varying bridge lengths. **(G)** Statistical tree representation of the experimental probabilities for elongated and compacted states over three generation of successive crossings. **(H)** Histogram representation of the probabilities for elongated, P(E), and compacted, P(C), states for various combinations of morphological switches: elongated to elongated P(E|E), compacted to compacted P(C|C), elongated to compacted P(E|C) and compacted to elongated P(C|E). Confocal microscopy images in super-resolution mode of side and top views of **(I)** elongated and **(J)** compacted cell morphologies. Actin is in grey and the nucleus in blue. The zoom of the top view in **(J)** shows the thickening of the actin cortex in compacted cells. Images were inverted. Scale bars, 20 µm for (I) and 5 µm for (J). **(K)** Plot profile of the normalized actin intensity in the actin cortex for elongated and compacted cells. The light purple zone indicates the distance corresponding to the full width at half maximum (FWHM) in compacted cells. **(L)** FWHM for elongated (orange, n=34) and compacted (purple, n=35) cells. Sequence showing typical cell morphology during successive crossings after treatment with **(M)** LatB and **(N)** Y27632. LatB or Y27632 were added in the culture media after the first crossing (CTRL). CTRL is DMSO for LatB and water for Y27632. Color-coded trajectories of cells treated with **(O)** LatB and **(P)** Y27632 at t=0 during their migration on a dumbbell pattern with a 160 µm long bridge. Elongated morphologies are in orange, compacted morphologies in purple, and square zones in grey. Mean crossing speed for cells before and after treatment with **(Q)** LatB (n=5; N=4) and **(R)** Y27632 (n=5; N=4). (S) Cell Shape Index (n=21 for CTRL in grey, n=28 for LatB in light blue and n=20 for Y27632 in light green; 2≤N≤4). **(T)** Successful crossing rate (n=146 for CTRL in grey, n=65 for LatB in light blue and n=58 for Y27632 in light green; 2≤N≤4). ****p < 0.0001 and n.s. not significant.

Previous theoretical work suggests that the polarity dynamics of cells adapt to local micropattern geometry(*24*, *25*), raising the question of whether a dependence of dynamical switching on local confinement could explain the dependence of morphological states on bridge length. Since a model with constant fixed switching rate cannot capture the length-dependence of compacted states **(Supp. Theory Note)**, we extended our model to include geometry-sensitive stochastic switching between elongated and compacted cells **(Fig. 3E and Movie S8)**. Specifically, we hypothesized that cells are more likely to transition to a polarized (*α* < 0), i.e. compacted, state when they were highly confined on the bridge, and more likely to transition back to an unpolarized (*α* > 0) state on the island. This is implemented by assuming that the switching rates *r*_E→C_(*x*) and *r*_C→E_(*x*) are position dependent and determined by the local width of the confinement **(Fig. 3E**, **Supp. Theory Note**). This simple model predicted an increase in the percentage of compacted morphologies with bridge length **(Fig. 3F)**, as observed experimentally **(Fig. 3D)**. Furthermore, the model predicted a further increase in the percentage of compacted cells in confining systems without islands (to 70%, **Supp. Fig. 2A**). To test this prediction, we confined cells to 1D micropatterned lines of 500 microns long **(Supp. Fig. 2B)** and we analyzed their shape during their back-and-forth motion **(Supp. Fig. 2C** and **Movie S10)**, indicating an increase of the percentage of compacted cells **(Supp. Fig. 2D)**, as predicted by the model. Together, these findings indicate that cells dynamically switch between elongated and compacted morphologies in a geometry-sensitive manner.

## Long-term memory links morphological states across transitions

We next investigated the overall time-scales of the switching process. A key parameter in our model is the average time-scale between switches, *T*. If this time-scale exceeds the average dwell time *τ*, the morphological state of cells in subsequent transitions is expected to be correlated. During any given transition, the probability of a cell being either elongated or compacted is 50% (*L*=160um, **Figs. 3G-H and Supp. Fig. 3**). In the absence of long-term memory (*T* << *τ*), we expect these probabilities to be independent of the morphological state during the previous transition **(Fig. 3H, Supp. Theory Note)**. In contrast, if *T* > *τ*, cells are more likely to remain in the same morphological state **(Fig. 3H)**. Experimentally, we indeed found strong correlations across transitions: following a compacted transition, 80% of cells in the subsequent transition were also compacted **(Figs. 3G-H)**. Conversely, 70% of previously elongated cells remained elongated in the next transition. These asymmetries are further enhanced in the third subsequent transition, to 90% and 80%, respectively **(Fig. 3G)**. These correlations across transitions are quantitatively predicted by our model for a switching time *T* ≈ 2.3*τ* ≈ 10 h. Together, these findings demonstrate that cells “memorize” their morphological state across transitions. This raises a key question: how is this memory is achieved, given that entry into the square is expected to lead to cell spreading and loss of polarization?

## Compacted cells have a thicker actin cortex

To identify the origin of the long-term memory of short-term confining events over short periods of time, we first hypothesized that migrating cells can remodel their matrix by secreting ECM proteins. We therefore investigated the possibility of physicochemical footprints left by migrating cells during migration, which could potentially result in history-dependence of migration behaviors over an extended period(*26*). We analyzed the biochemical composition of the FN micropattern after 5 hours of cell incubation and at the end of the 20-hour imaging period **(Supp. Figs. S4A)** using immunostaining for pre-coated and cell-produced fibronectin(*27*) **(Supp. Figs. S4B-C)**, as well as laminin **(Supp. Figs. S4D-E)**. Our findings indicated no statistical difference in fibronectin or laminin intensity between the different conditions, suggesting no direct relation to a biochemical remodeling of the surface.

We therefore hypothesized that the organization of the cytoskeleton could give rise to a mechanical memory across transitions. To verify this, we studied the organization of the cytoskeleton using super-resolution confocal microscopy, focusing on microtubules(*28*) and the actin cortex(*29*), which are heavily involved in the polarization of cells migrating in confined environments and in controlling changes in cell shape, respectively. Microtubules, which were evenly distributed in elongated cells and oriented along the bridge axis **(Supp. Fig. S5A)**, were significantly redistributed to the rear of compacted cells **(Supp. Fig. S5B)**. Interestingly, we observed a preferential orientation along the bridge axis at the front of both morphologies **(Supp. Fig. S5C)** and a more pronounced orientation along the vertical axis at the rear of compacted cells **(Supp. Fig. S5D).** The spatial redistribution of microtubules led to significant symmetry breaking in compacted cells **(Supp. Fig. S5E)**. Surprisingly, our observations of actin cortex organization in elongated **(Fig. 3I** and **Movie S6)** and compacted **(Fig. 3J** and **Movie S7)** morphologies indicated a doubling of the actin cortex thickness in compacted cells **(Figs. 3K-L)**. Recent studies have suggested that the thickness of the actin cortex is a fundamental aspect of regulating the internal stress and tension of the cortex, thus contributing to the shape of individual cells(*30*). Altogether, our results suggest that cortex thickening can serve as a mechanical memory of past confined events by maintaining a compacted shape.

## Mechanical memory is controlled by the actin cortex

Assuming that mechanical memory is associated with reinforced front-rear polarity, microtubules and the actin cortex are likely candidates to explain the origin of this mechanical memory. Indeed, it is well established that microtubules play a pivotal role in the establishment of cell polarity(*31–33*), while the cortex thickening can stabilize the cell compacted shape, generate long-range membrane tension propagation(*34*) and therefore contribute to the maintenance of the cell polarity(*35*). To test these hypotheses, we first treated cells with nocodazole, which disrupts microtubule dynamics by binding to tubulin and leading to microtubule depolymerization **(Supp. Fig. S6A** and **Movie S11)**. Tubulin-treated cells exhibited a similar CSI on the bridge to control cells **(Supp. Fig. S6B)**, a similar area on squares **(Supp. Fig. S6C)**, but a very low success crossing rate **(Supp. Fig. S6D)** and a very high dwell time **(Supp. Fig. S6E)**. Indeed, nocodazole-treated cells mostly became unable to migrate into the bridge and to maintain sufficient polarization to migrate within the confined area. Altogether, these results demonstrated that microtubules are not involved in the mechanical memory process related to maintaining the compacted shape but they are necessary for enabling the passage of epithelial cells in narrow environments.

In the next step, cells were treated either with a low concentration (20 nM) of Latrunculin B (LatB, **Fig. 3M** and **Movie S12**) to weaken cortical actin(*34*)b, or with Y27632 **(Fig. 3N** and **Movie S13)**, which inhibits Rho-associated protein kinase (ROCK), a key regulator of the actin cortical tension, leading to a significant decrease in cortex elasticity(*36*). Interestingly, cortical tension is higher in cells with elevated ROCK activity compared to those with higher Rac1 activity, suggesting that cortical tension increases when contractility dominates over actin polymerization(*37*). We quantified back-and-forth cell movements during an initial 4-hour control period, followed by an additional 20-hour period after adding one of the pharmacological agents **(Figs. 3O and P)**. Cells treated with LatB and Y27632 were significantly slower **(Figs. 3Q and R)**, exhibited a very low shape index on the bridge (**Fig. 3S**, CSI ≈0.2), characteristic of an extended morphology, and a much lower success rate of passage **(Fig 3T)**. These findings collectively demonstrate the major role of the actin cortex and the associated ROCK contraction pathway in maintaining the compacted shape.

## Perturbing the mechanical memory dynamics

Our finding that the morphological states of cells on the confining bridge are correlated across transitions suggests that cells can retain memory of previous states during the periods of unconfinement on the square islands. Indeed, the spreading rate on the square islands **(Supp. Fig. 7A-B)** can significantly affect the actin cortex thickness **(Supp. Fig. 7C)** and thus perturb the mechanical memory. This raises the question of whether the two morphological states lead to distinct cytoskeletal organization during the unconfining event on the islands. By studying transitions between two elongated (E/E, **Fig 4A**) or two compacted (C/C, **Fig 4B**) shapes, we observed that the cell area on squares of 1600 µm^2^ was statistically larger for E/E than for C/C **(Fig 4C)**, with a transition occurring at a cell area of around 1000 µm^2^. Additionally, we found that LatB- and Y27632-treated cells exhibited larger spreading areas **(Fig 4D)** and larger dwell times **(Fig. 4E)** on squares compared to compacted cells. Interestingly, we observed a continuous decreased of CSI **(Supp. Fig. 8A-B)** and crossing velocity **(Supp. Fig. 8C-D)** after each crossing for LatB- and Y27632 treated cells, while the spreading area on squares increased continuously in both conditions **(Supp. Fig. 8E-F).** Altogether, these results demonstrate that both the actin cytoskeleton and the ROCK contraction pathway control dwell times through cell spreading and actin cortex remodeling.

**Figure 4.**
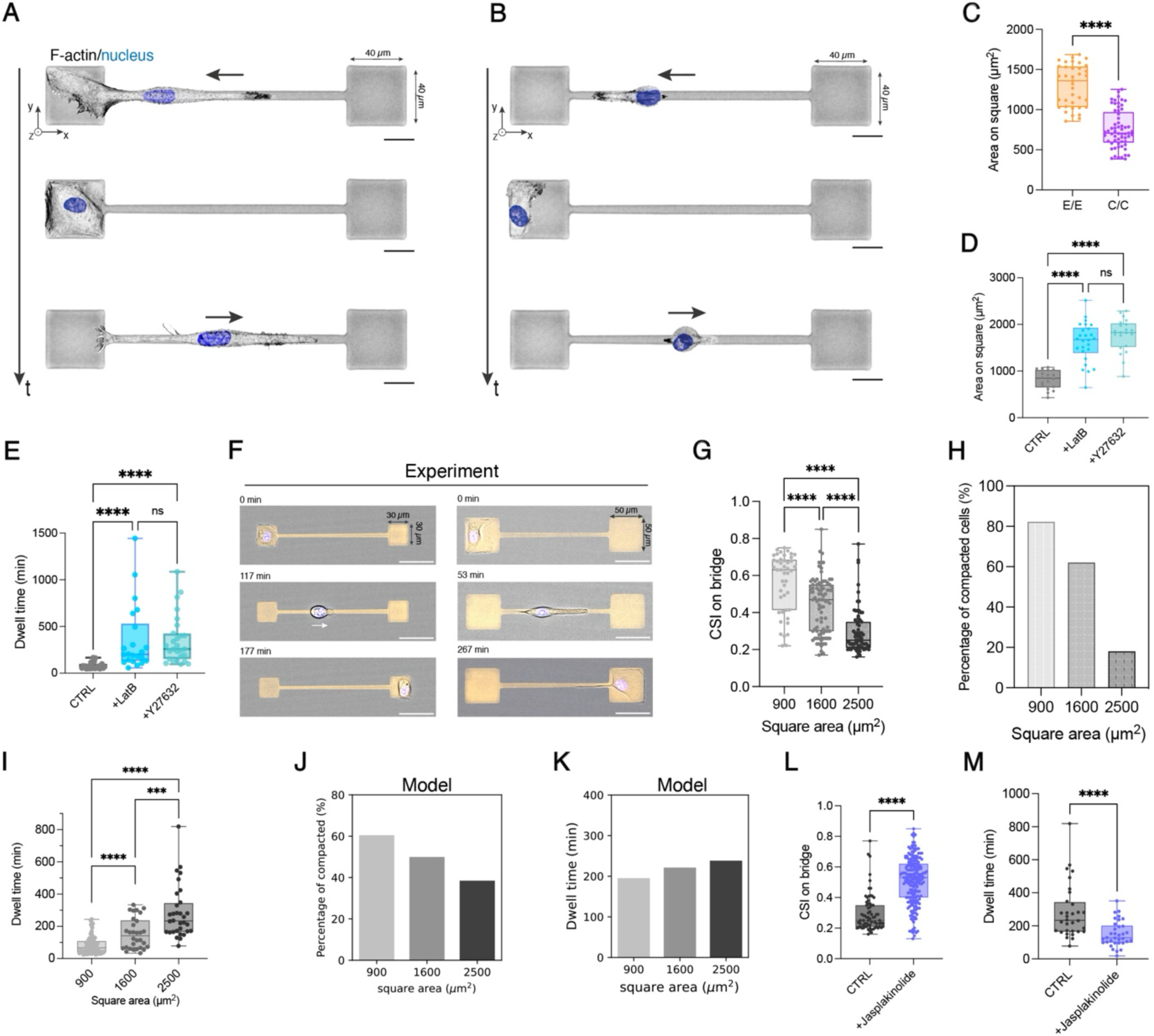
Perturbation of the mechanical memory dynamics. Representative three-step sequence showing a transition between **(A)** two elongated states (E/E) and **(B)** two compacted states (C/C). **(C)** Spreading area on 1600 µm² squares during a transition between two elongated states (E/E, n = 37 and N=3, orange) and two compacted states (C/C, n=61 and N=3, purple), indicating a transition occurring around 1000 µm². **(D)** Mean cell spreading area and **(E)** dwell time on 1600 µm² squares for control cells (n=27 and 2≤N≤ 4, grey), LatB-treated cells (n=22 and 2≤N≤ 4, light blue), and Y27632-treated cells (n=27 and 2≤N≤ 4, light green). **(F)** Representative three-step sequence showing a crossing event on a bridge of L=160 µm and W=6 µm connected to squares of 900 µm² (left) and 2500 µm² (right). **(G)** Cell shape index and **(H)** percentage of compacted cells (in purple) on a bridge 160 µm long and 6 µm wide for dumbbell geometries with squares of 900 µm² (n=45, N=3), 1600 µm² (N=87, N=14), and 2500 µm² (n=72, N=3). **(I)** Dwell time observed on squares of 900 µm² (n=45 and N=2), 1600 µm² (n=87 and N=16), and 2500 µm² (n=13 and N=3). Model predictions of **(J)** the percentage of compacted cells (in purple) on a bridge 160 µm long and 6 µm wide for dumbbell geometries with squares of 900 µm², 1600 µm², and 2500 µm² and **(K)** the dwell time on squares of 900 µm², 1600 µm², and 2500 µm². **(L)** Cell shape index (n=13 for CTRL in grey with N=4; and n=207 for Jasplakinolide in purple with N=4) and **(M)** dwell time for control cells (n=23 for CTRL in grey and N=4 for Jasplakinolide in purple, grey) and jasplakinolide-treated cells (n=34 and N=4, blue). CTRL is DMSO for Jasplakinolide experiments. **p < 0.01, ***p < 0.001, ****p < 0.0001; n.s. = not significant.

To test the role of available spreading area in the memory of morphological switching, we designed two new dumbbell geometries: a first one with two squares of 900 µm^2^, which is lower that the transition observed in **(Fig. 4C)**, and a second one with two squares of 2500 µm^2^, which are larger than the maximal spreading area **(Fig. 4D)**. Conceptually, squares of 900 µm^2^ will constrain cells into a compacted state, while larger squares of 2500 µm^2^ will provide the space for cells the opportunity to be in the elongated mode. By tracking back-and-forth motions on these new geometries **(Fig. 4F** and **Supp. Movies S14 and S15)**, we observed that the mean CSI on a bridge connected to 900 µm^2^ squares (0.56±0.16) was statistically larger than the mean CSI on a bridge connected to 1600 µm^2^ (0.44±0.15), while cells migrating on dumbbells with 2500 µm^2^ squares exhibited a lower CSI (0.29±0.12) **(Fig 4G)**, suggesting that the larger the square, the lower the CSI on the bridge **(Supp. Fig. 9A)**. Furthermore, our results indicate that the percentage of compacted cells was very high on bridges connected to 900 µm^2^ squares and decreased as the square area increased **(Fig. 4H)**. Additionally, increasing the square area leads to a proportional increase in dwell time **(Fig. 4I** and **Supp. Fig. 9B)**. We then expanded our model to reproduce the new dumbbell geometries and test its prediction in response to changes of square areas. As shown in **Figs. 4J** and **4K**, theoretical predictions confirmed that the larger the area of the square, the lower the proportion of compacted cells **(Fig. 4J)** and the longer the dwell time **(Fig. 4K)**, demonstrating the validity of our model. Altogether, these results demonstrate that modulating the dwell time allows tuning the mechanical memory of confined cells.

To demonstrate the crucial role of the actin cortex in maintaining cell shape memory, we treated cells migrating on dumbbells connected to a deconfinement square size of 2500 µm² with jasplakinolide (Jas, **Movie s16**), a pharmacological agent that stabilizes actin filaments(*38*). We used a moderate concentration of Jas, which has been shown to stabilize actin filaments and increase the actin cortex thickness(*39*), without significantly impacting the recycling of actin filaments in the lamellipodium, which is necessary for cell migration(*40*). Our results indicated that cells treated with Jas continued to migrate, adopted a more compacted shape (CSI=0.51±0.13) **(Fig. 4L)** and exhibited a significantly shorter residence time (τ=151±67 min) **(Fig. 4M)** than control cells. These results demonstrate that the reinforcement of the actin cytoskeleton enables cells to maintain a compact shape even when placed in very large deconfinement areas that would typically normally lead to elongated shapes **(Fig. 4F)** and high residence times **(Fig. 4H)**.

## Discussion

Collectively, our data identify a morphological switch in confined migration and establish a mechanical memory function of the actin cortex. The dynamic switching between compacted and elongated morphologies allows cells to alternate between highly polarized, directed and more exploratory, undirected motility. This adaptability likely plays a key role in navigating through heterogeneous matrices and narrow constrictions, relevant to various physiological processes, such as immune cell patrolling, progenitor cell motility during development, and cancer cell invasion. The mechanical memory we describe allows cells to retain a record of their previous morphological state, even as they transition through subsequent cycles of confinement and unconfinement. This memory is encoded by the organization of the actin cortex, enabling cells to maintain a compacted during temporary unconfinement events. Our findings align with previous reports that cell spreading regulates cortex thickness(*41*) and that strengthening the actin cortex facilitates cell migration(*42*, *43*). The retention of memory from previous confining events may allow cells to traverse heterogenous confinements such as interstitial spaces without having to pause to reorganize their shape each time they encounter an unconfined space. Notably this mechanical memory preserves a compacted, highly polarized state, promoting rapid exploration through persistent motion and efficient navigation at dead ends.

From a theoretical perspective, our findings highlight the complex, time- and geometry-dependent dynamics of the intrinsic active self-propulsion of migrating cells, challenging the conventional assumption that this parameter is constant in active matter systems(*23*). The observed morphological switch and its associated memory dynamics may also have significant implications for collective cell behavior in confined environments, such as flocking transitions governed by the polarization state of cells(*44*). While our focus here has been on identifying a minimal model to predict the long time-scale statistics of morphological switches, future theoretical work will be needed to link these dynamics to mechanistic models of actin and polarity regulation(*25*, *45*, *46*).

To further elucidate the complex nature of this mechanical memory, key questions remain: how long must cells be confined for memory to be established? What are the changes in the 3D nanostructure of the actin cortex that determine the memory’s time-scale?(*47*) Taken together, our findings represent among the most direct evidence to date that morphological switches enhance efficiency during confined migration and support the role of the actin cortex as a mechanoadaptive mechanism involved in maintaining a cell memory of past morphology during confined invasion.

## Supporting information

Supplementary Theory Note

## Acknowledgements

The authors are grateful to members of the SG lab for feedback and suggestions. We thank Edouard Hannezo and Joachim O. Rädler for inspiring discussions. We thank Daniel Selma Herrador and Martial Balland for their help to improve the microprinting method. D.B.B. was supported by the NOMIS foundation as a NOMIS Fellow and by an EMBO Postdoctoral Fellowship (ALTF 343-2022). Y.K., M.L., and S.G. acknowledge funding from the University of Mons ‒ UMONS, FEDER Prostem Research Project no. 1510614 (Wallonia DG06), the F.R.S.-FNRS Epiforce Project no. T.0092.21, the F.R.S.-FNRS Cellsqueezer Project no. J.0061.23, the F.R.S.-FNRS Optopattern Project no. U.NO26.22 and the Interreg MAT(T)ISSE project, which is financially supported by Interreg France-Wallonie-Vlaanderen (Fonds Européen de Développement Régional, FEDER-ERDF). Y.K and M.L. are financially supported by F.R.S.-FNRS as FRIA Grantee FNRS and Postdoctoral Fellow (Chargé de Recherches), respectively. Y.K. and S.G. acknowledge le Fonds pour la Recherche Médicale dans le Hainaut (FRMH). GC was supported by a grant from the Biotechnology and Biological Sciences Research Council (BBSRC, BB/V007483/1).

## Author contributions

S.G. and Y.K. conceived the project and S.G. supervised the project. Y.K. developed micropatterns, performed time-lapse cell experiments, cell tracking and confocal imaging. D.B.B. developed the theoretical model and performed simulations and analysis. M.L. contributed to experiments and commented the manuscript. G.C. provided cell lines and commented the manuscript. Y.K., D.B.B. and S.G. analyzed data and prepared the figures. The article was written by Y.K., D.B.B. and S.G, read and corrected by all authors, who all contributed to the interpretation of the results.

## Competing interests

The authors declare no competing interests.

## Data availability

All data are available from the corresponding authors upon request.

## Code availability

Codes are available from the corresponding authors upon request.

## Materials and Methods

### Cell culture

MCF-10A cells were cultured in DMEM/F-12 (ThermoFisher) supplemented with 5% Horse Serum (ThermoFisher), 20 ng/mL epidermal growth factor (EGF) (Peprotech), 0.5 mg/mL Hydrocortisone (Merck), 100 ng/mL Cholera Toxin (Enzo Life Sciences), 10 μg/mL insulin from bovine pancreas (Merck) and 0.1% Penicillin/Streptomycin (Pen/Strep, Merck). Cells were passaged every 3-4 days when reaching confluence and plated at a 1:4 dilution (∼2 million cells/T75 flask). For cell passaging, the culture media was aspirated, and cells were washed with 10mL of 1x PBS. After aspirating the PBS, cells were incubated with 2mL of 0.25% trypsin (Merck) at 37°Cfor 10 minutes. Trypsinization was halted by adding 4mL of media, and cells were spun at 1.3 rpm for 5 minutes in a 15mL falcon tube to obtain a pellet. The old media was aspirated, and cells were resuspended in 1mL of fresh media before being plated. MCF-10A cells were used up to a maximum of 15 passages.

### Immunocytochemistry and live staining

MCF-10A epithelial cells were fixed with 4% paraformaldehyde (PFA) in PBS for 15 min at room temperature after cell migration experiments. Subsequently, cells were washed three times with PBS, with the last wash lasting for 5 min, and permeabilized using 0.05% Triton X-100 in PBS for 15 min at room temperature. After another round of three washes in PBS, permeabilized cells were blocked with a solution of 5 v/v% Fetal Bovine Serum (FBS, Gibco) and 1 w/v % Bovine Serum Albumin (BSA, Merck) in PBS for 30 minutes at room temperature. Actin filaments were stained with AlexaFluor 488 phalloidin (Invitrogen, 1:200), the nucleus with 4′,6-diamidino-2-phenylindole (DAPI; Invitrogen, 1:200), microtubules with an anti-tubulin antibody produced in mouse (Sigma-Aldrich, 1:200).

### Confocal microscopy

Images were collected in confocal mode with a Nikon A1R HD25 motorized inverted microscope equipped with ×20, ×40, ×60 Plan Apo (NA 1.45, oil immersion) and ×100 Plan Apo silicone objectives and lasers that span the violet (405 and 440 nm), blue (457, 477, and 488 nm), green (514 and 543 nm), yellow-orange (568 and 594 nm), and red (633 and 647 nm) spectral regions. Confocal images were recorded with ×100 Plan Apo silicone objective of high numerical aperture (Plan Apochromat Lambda S ×100 Silicone, Nikon Inc) in galvanometric mode with small Z-depth increments (0.1 μm) and a pinhole of 12 µm to capture high resolution images. Confocal images were recorded and processed using NIS-Elements (Nikon, Advanced Research version 4.5).

### Super-resolution confocal microscopy

The actin cytoskeleton and microtubule networks of MCF-10A cells were observed using confocal microscopy in super-resolution mode. Images were captured using a Nikon AX Ti2 Confocal microscope (Nikon, Japan) combined with the Nikon Spatial Array Confocal (NSPARC) detector. Z-stacks images were collected with a Galvano scanner using a 60x/1.42 Plan Apo oil immersion objective and a step size of 0.17 µm for three channels (DAPI, TRITC and FITC). Confocal images were processed using the NIS-Elements software (Nikon, Advanced Research version 4.5, Japan), with a Richardson-Lucy deconvolution method applied to remove unfocused signal.

### Time-lapse imaging

Forth and back motions on dumbbell micropatterns were recorded at ×40 magnification with a photometrics Prime 95B camera (Photometrics Tucson, AZ) mounted on a Nikon A1R HD25 Ti2 motorized inverted microscope (Nikon, Japan). An incubation chamber (Okolab, Italy) was used to maintain CO2 levels at 5% and temperature at 37°C throughout the imaging session. Time-lapse images were recorded for 20 hours in DIC mode with a time interval of 3 min and processed using NIS-Elements (Nikon, Advanced Research version 4.5, Japan).

### Actin flow experiments

MCF-10A cells were incubated with SPY555-FastAct™ (Spirochrome) for 2 hours at 37°C following the manufacturer’s protocol (dilution 1:1000 in culture medium). Live actin imaging was performed in confocal mode on a Nikon A1R HD25 Ti2 motorized inverted microscope equipped with a Plan Apo 100x/1.35 silicone objective. An incubation chamber (Okolab, Italy) was used to maintain CO2 levels at 5% and temperature at 37 °C throughout the imaging session. Images were acquired at a rate of one 1.25 second for a total duration of five minutes. High-speed confocal images were recorded and processed using NIS-Elements (Nikon, Advanced Research version 4.5, Japan).

### Retrograde actin flow determination

To determine the retrograde actin flow at the cell leading edge (lamellipodium), time-lapses were initially converted to 8-bit format, underwent background subtraction, contrast enhanced, and were subjected to a Gaussian blur filter (sigma=1.5). Subsequently, kymographs were generated using the reslice command tool on Fiji. Kymograph were obtained from 1 pixel wide lines drawned at the center of the lammelipodia. Three lines were drawn per lamellipodia and the XY coordinates of the line was saved to calculate the slope corresponding to the flow speed value in units of µm/s, which is subsequently converted to nm/s.

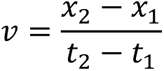

### Protein micropatterning

The micropatterning of fibronectin on glass was performed according to the light-induced molecular adsorption of protein method using the PRIMO system (Alvéole, France) mounted on an inverted microscope (Nikon Eclipse Ti2, Japan), as previously described(*48*, *49*). Briefly, glass bottom dishes (Cellvis) and PDMS stencils are plasma-activated for 3 min (Harrick Pasma), prior to their assembly. The stencils allow to minimize volume solution and avoid fast dewetting of the surface. The glass surface was coated using a 0.1% (m/v) poly-L-lysin for 30 min at room temperature and incubated 1 hour at room temperature with a 100 mg/ml PEG-SVA passivation solution (polyethylene glycol-Succinimidyl Valerate). After drying, the surface is covered with a mixture of PLLP-gel photoinitiator solution (Alvéole, France), surfactant and 70% ethanol and leave to dry 30 min before being transferred to the microscope stage. Designed motifs are projected onto glass through the Primo Digital Micromirror Device (DMD) using Leonardo software (Alvéole, France). A dose of 100 mJ/mm^2^ at 375 nm UV light illuminating (100% laser power) focused through a 20x objective (Nikon, Plan Fluor) was used to activate the photoinitiator and remove the passivation layer at desired locations. The surface is washed and coated with 15 µg/ml human plasma fibronectin (Sigma) containing 5 µg/ml of fluorescent human plasma fibronectin for 5 min at room temperature. Patterned surfaces can either be used directly or stored at 4°C in 0.1 M NaHCO3 buffer (pH=8.3) up to 24 hours after preparation.

### Drug treatments

Actin depolymerization, ROCK inhibition and microtubule depolymerization treatment were achieved upon addition of 20 nM Latrunculin B, 10 µM Y27632 and 2.5 µM nocodazole, respectively, to samples after four hours of timelapse imaging. Then imaging was resumed for an additional twenty hours in presence of the drug. Actin stabilization treatments were performed by incubating samples with 1 nM of Japlakinolide 30 min before the beginning of the timelapse experiments. LatB, nocodazole, and jasplakinolide were diluted in DMSO, while Y27632 was diluted in water.

### Cell tracking

Time-lapse sequences were recorded for 20 hours in DIC mode with a time interval of 3 min, leading to approximately 400 frames per experiments. These sequences were imported as a stack in Fiji, converted to 8-bit format, and analysed using Cellpose 2.0(*50*), a deep learning-based cell segmentation, that provides excellent segmentation of cell outlines in DIC and phase contrast images. Cellpose was incorporated into TrackMate(*51*), a user-friendly tracking plugin for Fiji. Cell outlines obtained for each frame manually verified and then analyzed to determine the cell spreading area, the cell perimeter and the coordinates of the cell front, cell rear, and center of mass.

### Cell shape index (CSI)

Areas (A) and perimeters (P) of migrating cells were measured with ImageJ on DIC images using CellPose 2.0. Both geometrical parameters were used to calculate the CSI from the relationship:

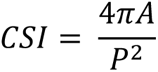

The CSI assumes values between one (circular shape) and zero (elongated, linear morphology).

### Physicochemical footprints

Given that the physicochemical footprints left by migrating cells over an extended period can modulate their migration speed and trajectory(*26*), we analyzed the biochemical composition of the FN micropattern after 5 hours of cell incubation and at the end of the 20-hour imaging period **(Fig. S1A)**. Samples were fixed at both short (t=5 hours) and long (t=20 hours) experimental times and immunostained for immunostained for pre-coated and cell-produced laminins (Sigma-Aldrich) and fibronectins, using an antibody specific to the fibronectin produced by the cells themselves (Fibronectin IST-9, Santa cruz biotechnology) (*27*).

### Statistical analysis

Each experiment was repeated at least three times. Every set of data was tested for normality test using the d’Agostino-Pearson test in Prism 10.0 (GraphPad) that combines skewness and kurtosis tests to test whether the shape of the data distribution was similar to the shape of a normal distribution. For paired comparisons, significances were calculated in Prism 10.0 (GraphPad) with a Student’s *t*-test (two-tailed, unequal variances) when the distributions proved to be normal. If a data set did not pass the normality tests, the significances were calculated with Mann–Whitney (two-tailed, unequal variances). For multiple comparisons with non-normal distribution, data sets were analyzed with a Kruskal-Wallis test in Prism 10.0 (GraphPad), which is a suitable nonparametric test for comparing multiple independent groups when the data are skewed. When the null hypothesis was not retained (p-value < 0.05), Kruskal-Wallis was corrected with Dunn’s test, which is a nonparametric test with no pairing and multiple comparisons that can be used for both equal and unequal sample sizes. Unless otherwise stated, all data are presented as mean ± standard deviation (s.d.). The confidence interval in all experiments was 95% and as a detailed description of statistical parameters it is included in all figure captions with *p < 0.05, **p < 0.01, ***p < 0.001, ****p<0.0001 and n.s. is not significant.

**Supplementary Figure 1.**
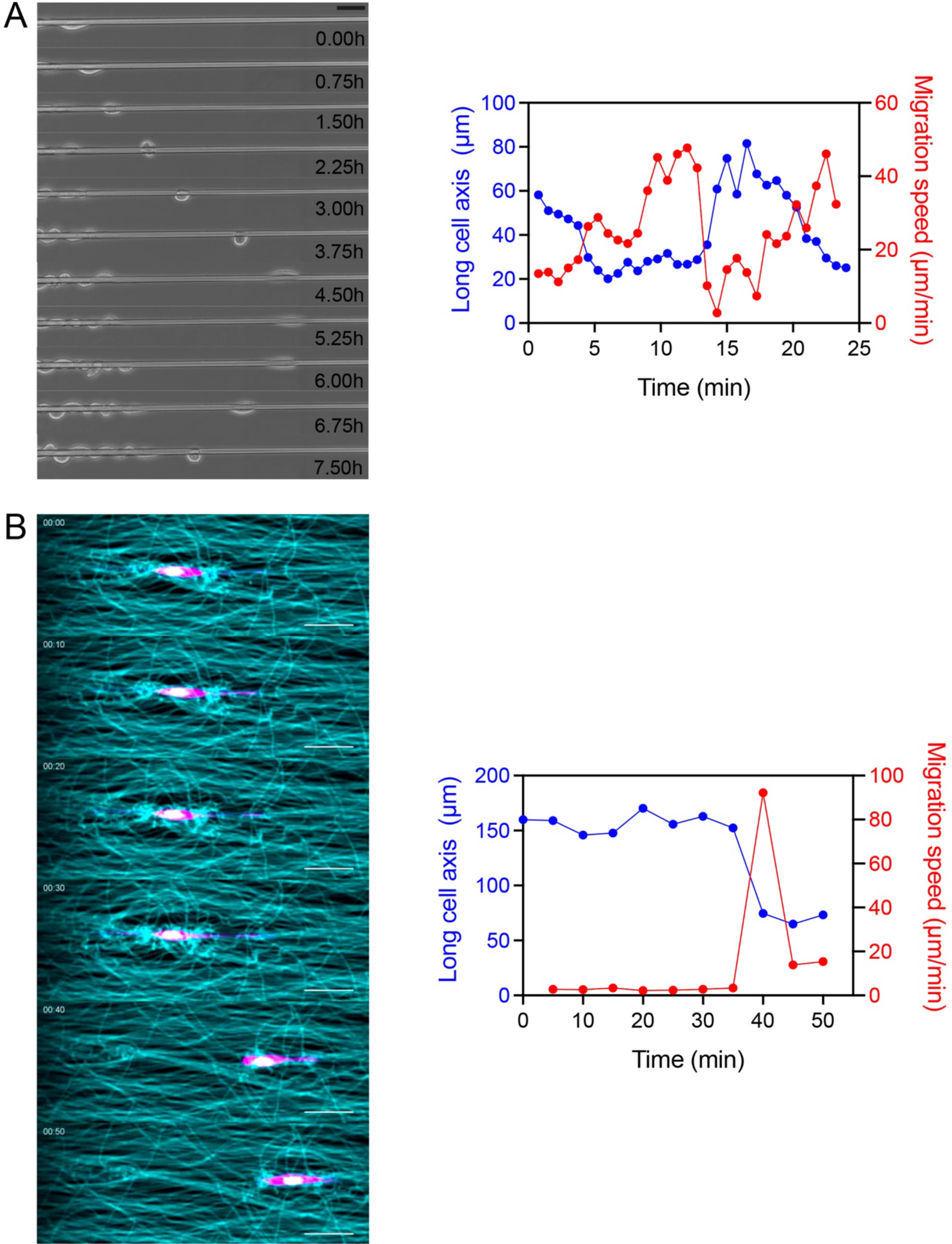
Examples of morphological switch on glass fiber and in 3D collagen fiber matrix. **(A)** Time-lapse sequence of a single MDCK epithelial cell migrating on a smooth glass wire (diameter = 5.3 µm). After detaching from the tissue, the epithelial cell rounds up and migrate on the glass fiber. The temporal evolution of the long cell axis (blue) and the migration speed (red) indicates that the cell morphological switch from an elongated to a compacted morphology is associated with an increase in migration speed. Scale bar, 50 µm. Adapted with permission from (*20*). **(B)** Time-lapse sequence of a single fibroblast (NIH3T3, magenta) migrating in a 3D matrix composed of aligned collagen fibers (cyan). Scale bar, 50 µm. The temporal evolution of the long cell axis (blue) and the migration speed (red) indicates that the cell morphological switch from an elongated to a compacted morphology is associated with an increase in migration speed with a maximum speed around 80 µm/min. Adapted with permission from (*21*).

**Supplementary Figure 2.**
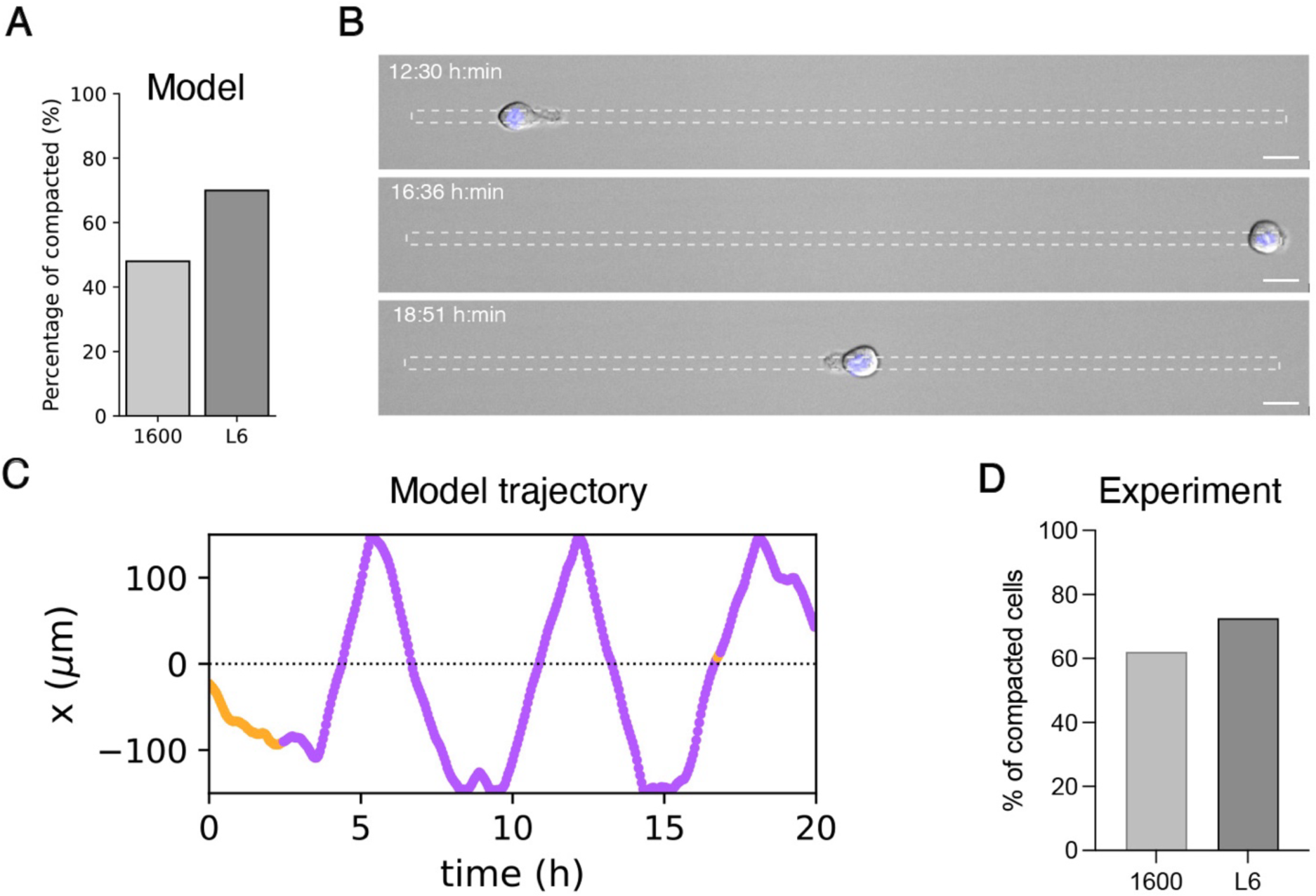
Semi-infinite narrow segments. **(A)** Time-lapse sequence of a compacted cell morphology migrating on a one-dimensional (1D) micropatterned line of W=6 µm and L=500 µm. Scale bar, 20 µm. **(B)** Percentage of compacted cells on the bridge of dumbbells (W=6 µm and L=160 µm) with square area of 1600 µm^2^ (n= 56, N = 16) versus 1D semi-infinite line of W=6 µm and L=500 µm (n=10, N = 2). **(C)** Representative color-coded trajectories: elongated morphologies in orange (CSI<0.4) and compacted morphologies in purple (CSI>0.4). **(D)** Percentage of compacted cell predicted by the model with elongated morphology in orange and compacted morphology in purple. On 6 µm line, percentage of compacted cell is expressed as the percentage of time spent under the compacted morphology over a 20-hour time-lapse.

**Supplementary Figure 3.**
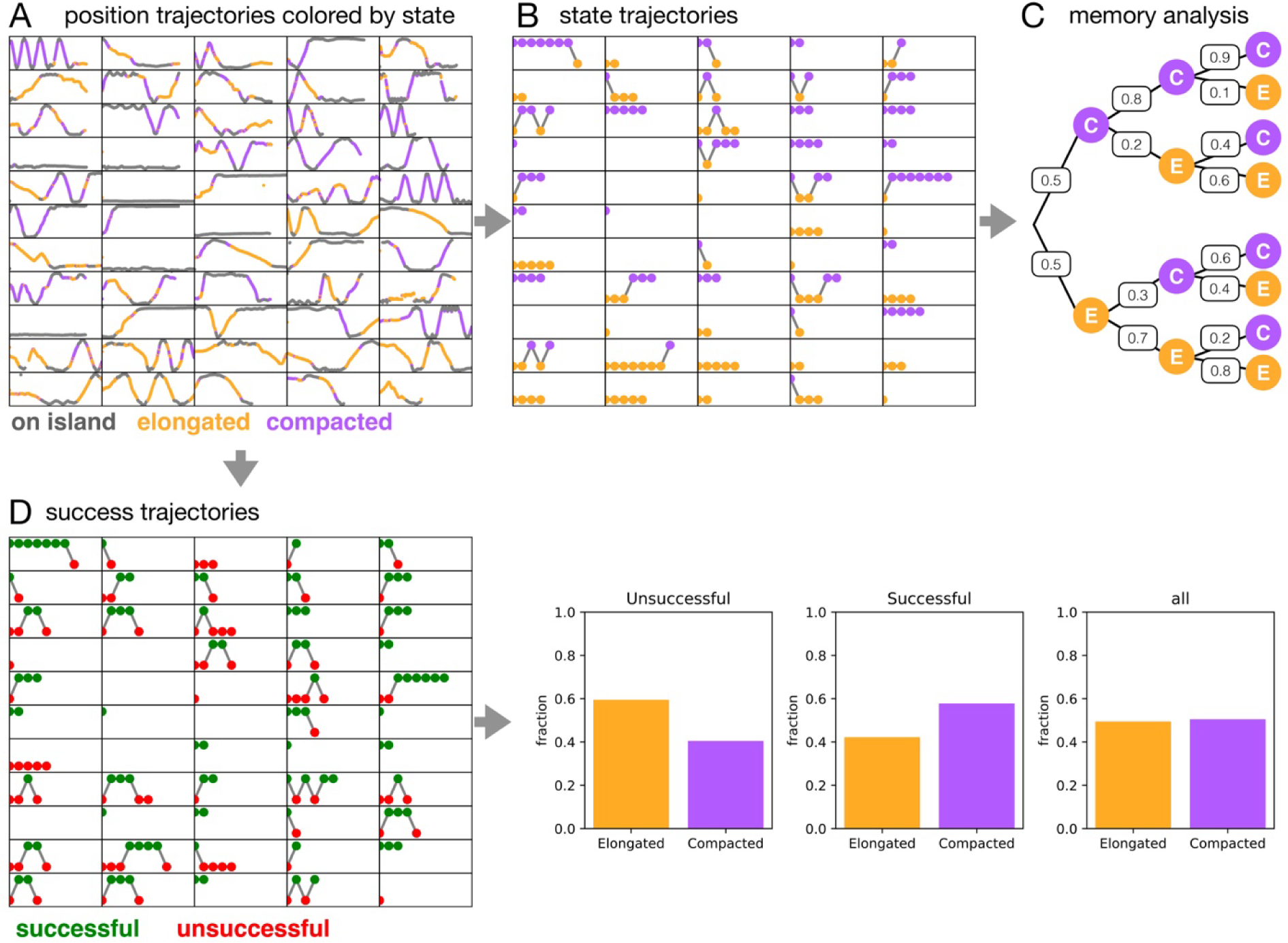
Analysis of memory dynamics from cell trajectories. **(A)** Selection of n=55 color-coded single cell trajectories of individual MCF-10A cells migrating on FN dumbbell micropatterns for 20 hours. Trajectories on the bridge are color-coded by state: elongated morphologies in orange (CSI<0.4) and compacted morphologies in purple (CSI>0.4), with square zones in grey. **(B)** State trajectories computed from position trajectories in (A). A morphological state is assigned to each transition on the bridge, based on the morphological state adopted for the majority of time-points during each transition. **(C)** Memory analysis tree diagram for events C (compacted transition) and E (elongated transition) obtained from state trajectories in B. Numbers of the branches indicate probabilities. **(D)** Trajectories of successful (green) and unsuccessful (red) transitions, defined as follows: in successful transitions, the cell (nucleus) enters the bridge and transmigrates to the other island. In unsuccessful transitions, the nucleus enters the bridge and returns to the same island. Fraction of unsuccessful, successful and total transition, which are repartitioned into two categories, elongated (orange) and compacted (purple).

**Supplementary Figure 4.**
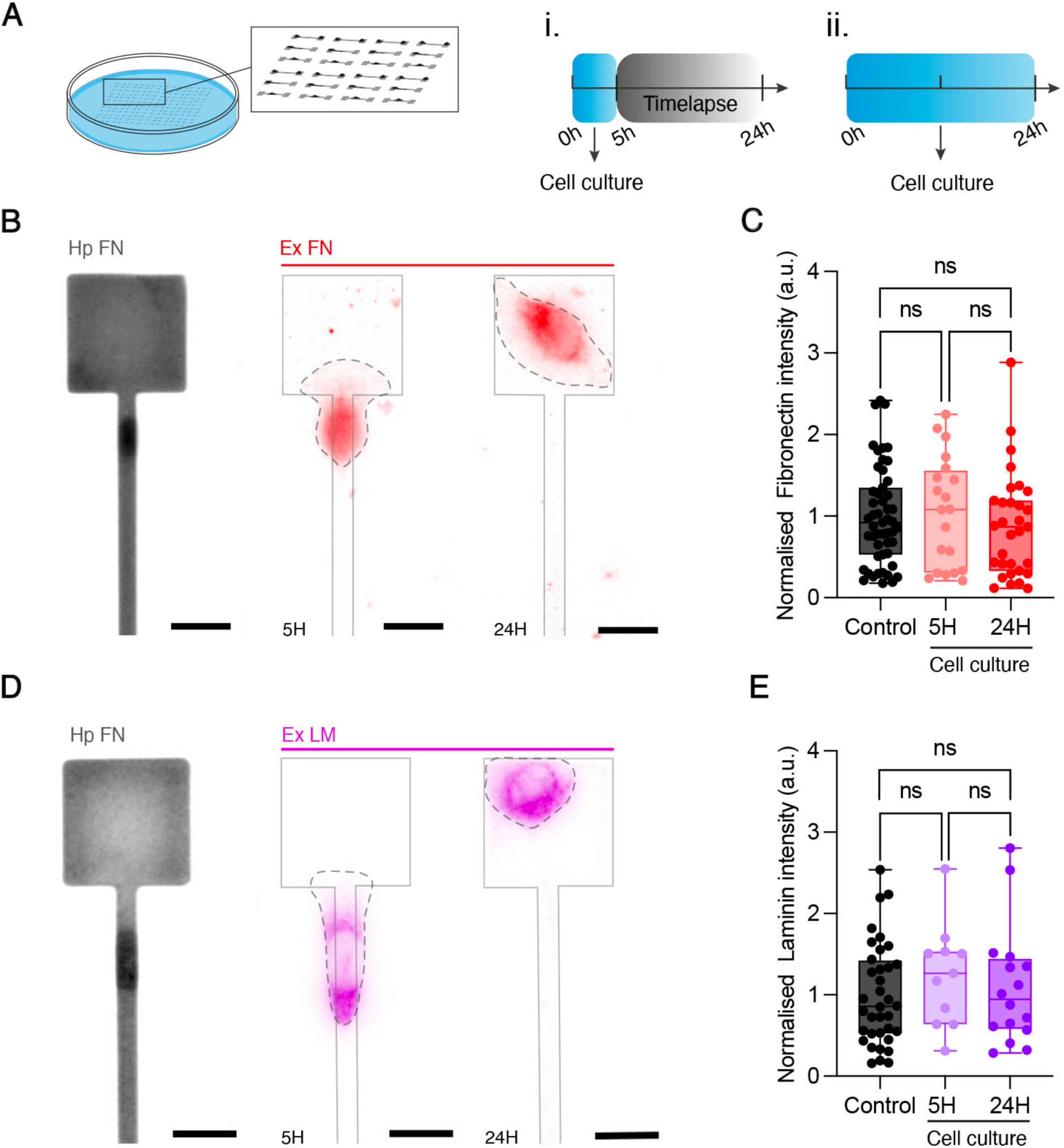
Physico-chemical footprints. **(A)** Schematic representation of the two experimental procedures utilized to determine pattern conditioning by migrating cells: cells are seeded on human plasma (hp) fibronectin (FN) micropattern and fixed either after (i) 5 hours (before the time-lapse) or (ii) 24 hours (after the time-lapse) of culture on micropatterns. **(B)** Epifluorescence images of hp FN (in grey) and cellular FN (in red) after t=5h and t=24h of cell culture. Scale bar, 20 µm. **(C)** Normalized FN intensity for control micropattern in black (n=51 patterns, N=2), 5 hours of conditioning in light red (n = 20 patterns, N=2) and 24 hours of conditioning in red (n=31 patterns, N=2). **(D)** Epifluorescence images of hp FN (in grey) and cellular laminin (LM, in purple) after t=5h and t=24h of cell culture. Scale bar, 20 µm. **(E)** Normalized LM intensity from control in black (n = 36 patterns, N = 2), 5 hours conditioning in light purple (n=11 patterns, N=2) and 24 hours conditioning in purple (n=16 patterns, N=2). n.s. = not significant.

**Supplementary Figure 5.**
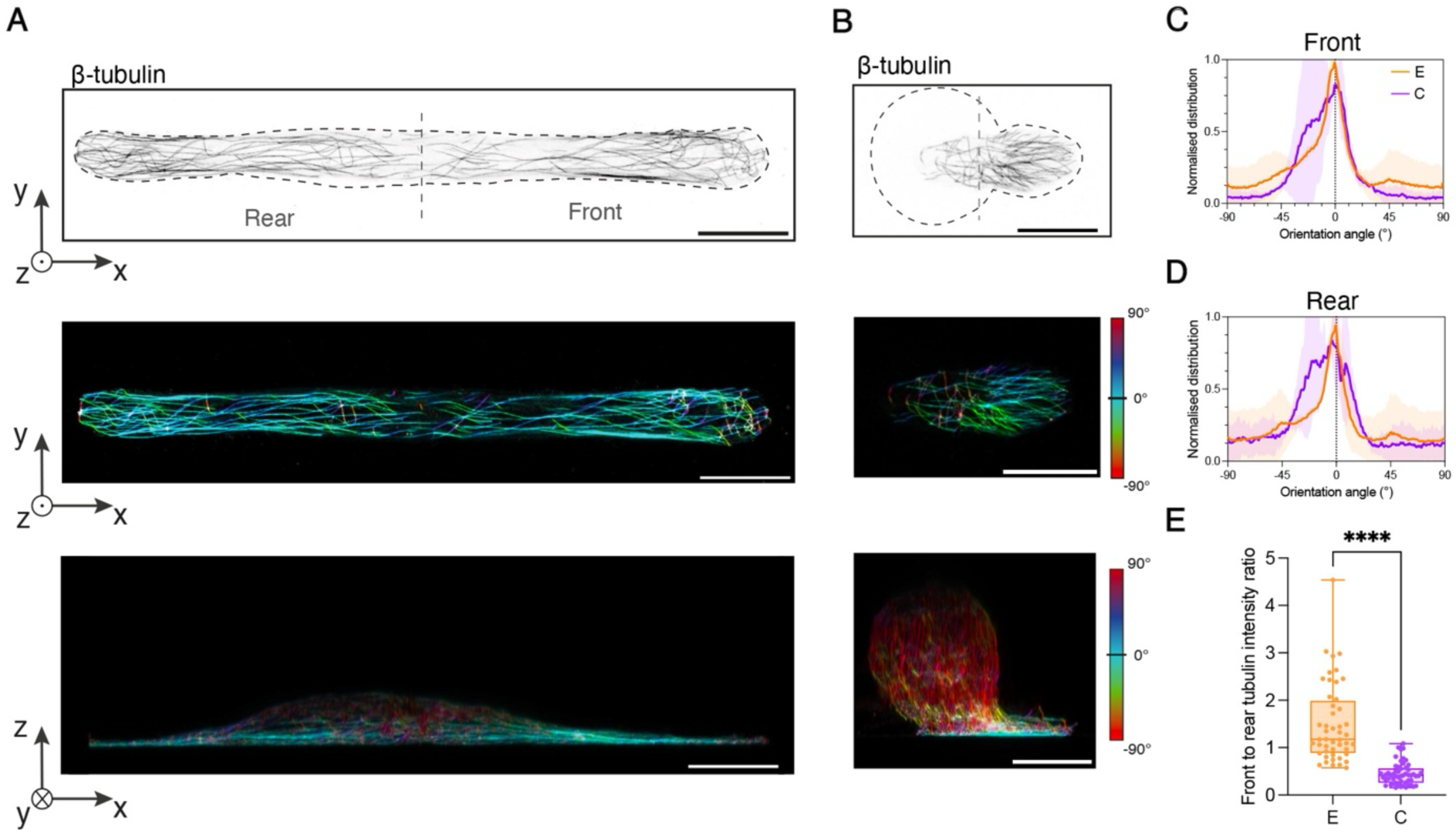
Spatial distribution of microtubule filaments. **(A)** Confocal microscopy images in super-resolution mode of the basal planes of elongated (left) and compacted (right) cell morphologies. β-tubulin staining is in dark grey (inverted image) and the cell body outline is represented with a black dashed line. Scale bar, 20 µm. **(B)** Front-to-rear tubulin intensity ratio of elongated (n=48, N=3, orange) and compacted (n=54, N=3, purple) morphologies during crossing events. **(C)** Confocal microscopy images in super-resolution mode of basal planes of elongated (left) and compacted (right) cell morphologies showing β-tubulin filaments color-coded according to their spatial orientation. Scale bar, 20 µm. Distribution of the normalized intensity of microtubules **(D)** at the front and **(E)** at the rear of elongated (orange, n=3, N = 2) and compacted (purple, n=3, N = 2) cell morphologies. **(F)** Confocal microscopy images in super-resolution mode of the basal planes of elongated (left) and compacted (right) cell morphologies showing the 3D projection side view of β-tubulin filaments color-coded according to their spatial orientation. Scale bar, 20 µm.

**Supplementary Figure 6.**
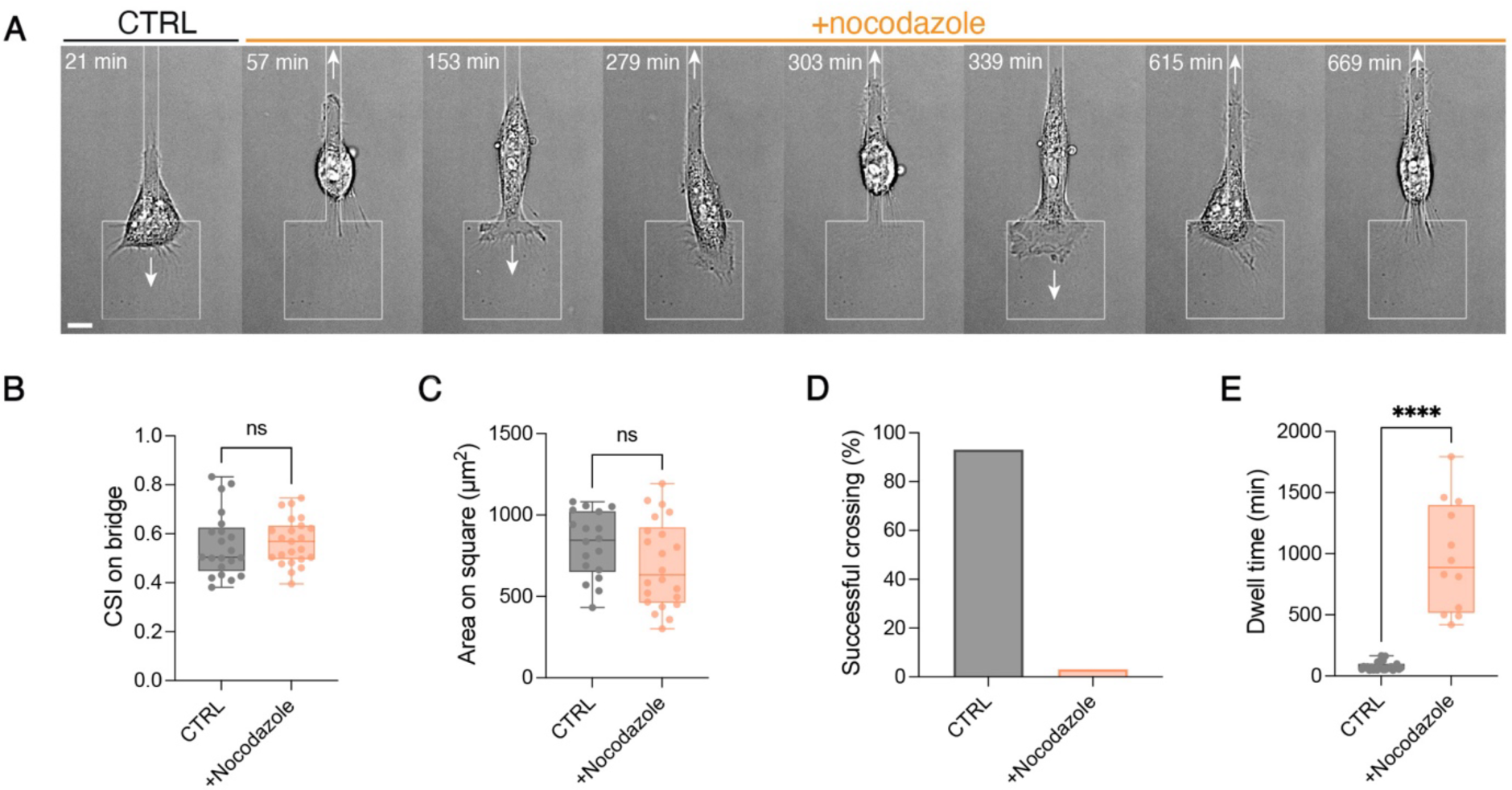
Effect of a nocodazole treatment on confined migration. **(A)** Time-lapse sequence of a compacted cell entering a bridge before (control, CTRL) and after treatment with nocodazole (orange). CTRL is DMSO for nocodazole experiments. Scale bar, 10 µm. White arrows show the direction of migration. **(B)** Cell shape index on bridge for CTRL (n=21, grey) and nocodazole-treated cells (n = 23, light orange). **(C)** Cell area on square for CTRL (n=18, grey) and nocodazole-treated cells (n=22, light orange). **(D)** Successful crossing percentage for CTRL (n=146, grey) and nocodazole-treated cells (n=33, light orange) and **(E)** dwell time on square for CTRL (n=22, grey) and nocodazole-treated cells (n=12, light orange). ****p < 0.0001; n.s. = not significant.

**Supplementary Figure 7.**
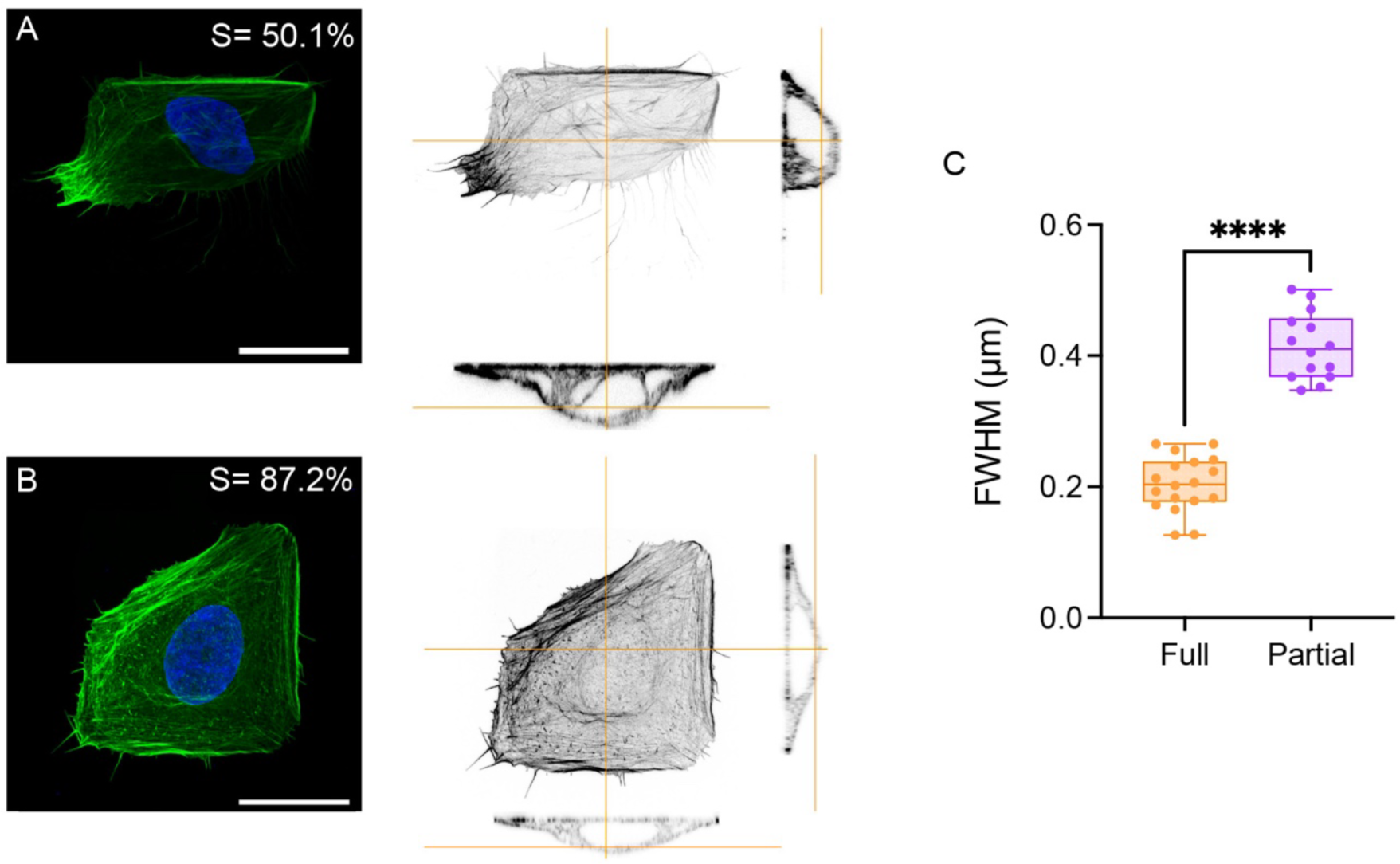
Thickness of the actin cortex in cells spread on squares. Super-resolution confocal images of **(A)** a partial spreading and **(B)** a full spreading of epithelial cells on FN square of 1600 µm^2^. Cells are immunostained for F-actin with phalloidin (in green) and DNA with DAPI (in bue). Inverted images show normal and side views of the actin cytoskeleton at the apical side. The spreading rate (S) is 50.1% for partial spreading and 87.2% for full spreading. Scale bars, 20 µm. **(C)** Full width at half maximum (FWHM) represents the actin cortex thickness for partial (in orange, n=18) and full (in purple, n=18) spreading conditions, with ****p < 0.0001.

**Supplementary Figure 8.**
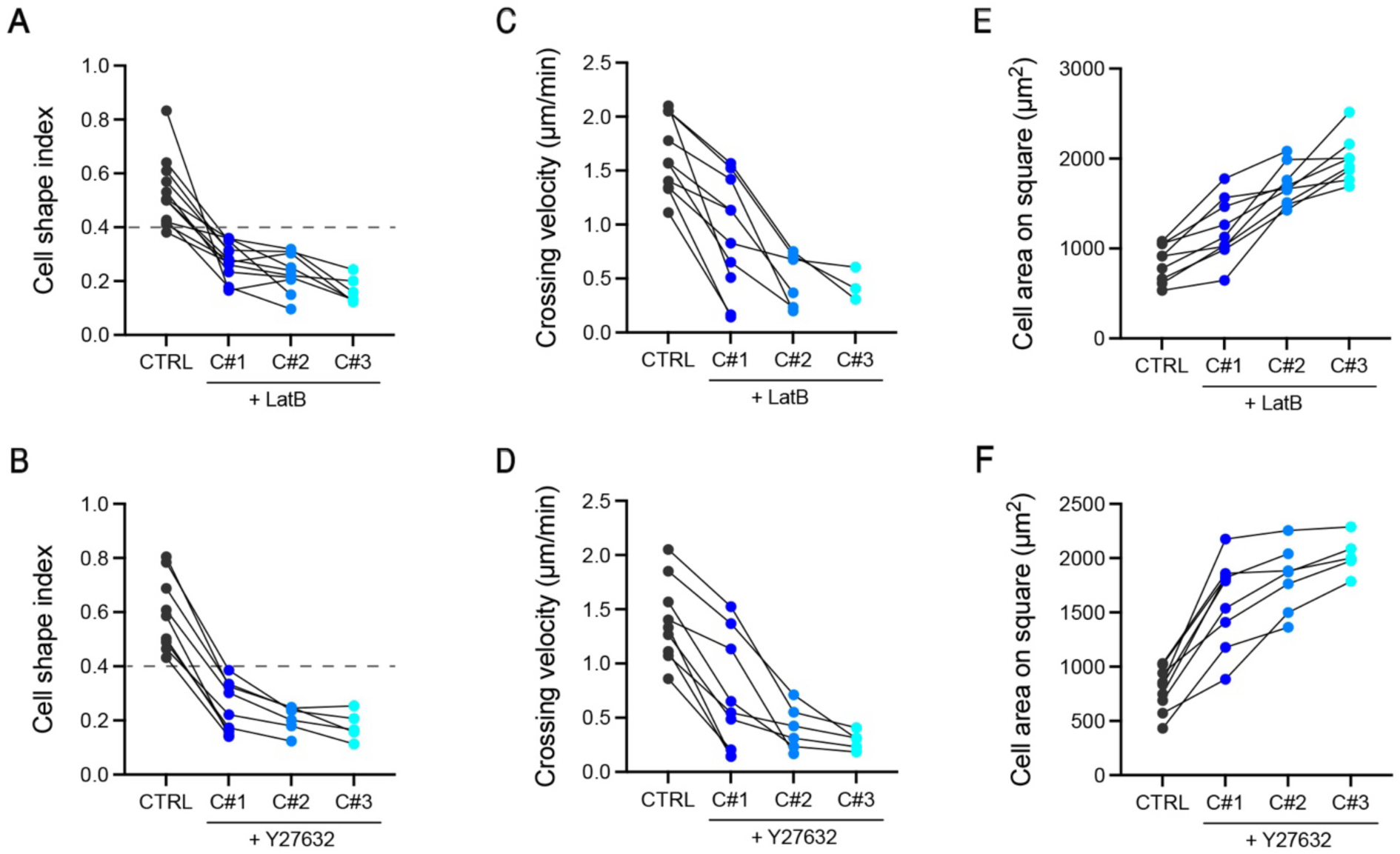
Evolution of the cell shape index, the crossing velocity and the cell area on square in response to Latrunculin B and Y27632 treatments. **(A)** Cell shape index (n=11, N=4), **(B)** crossing speed (n = 10, N = 4), and **(C)** and cell area on square (n = 9, N = 4) before (control, CTRL) and after treatment with latrunculin B (shades of blue). **(D)** Cell shape index (n=8, N=2), **(E)** crossing speed (n=9, N=4), and (F) cell area on square (n=9, N=2) before (control, CTRL) and after treatment with Y-27632 (shades of blue). Data from the same cell are connected by black lines. C#1, C#2, C#3 refer to the transition number after addition of the drug. Control experiments (CTRL) were performed in DMSO for both conditions.

**Supplementary Figure 9.**
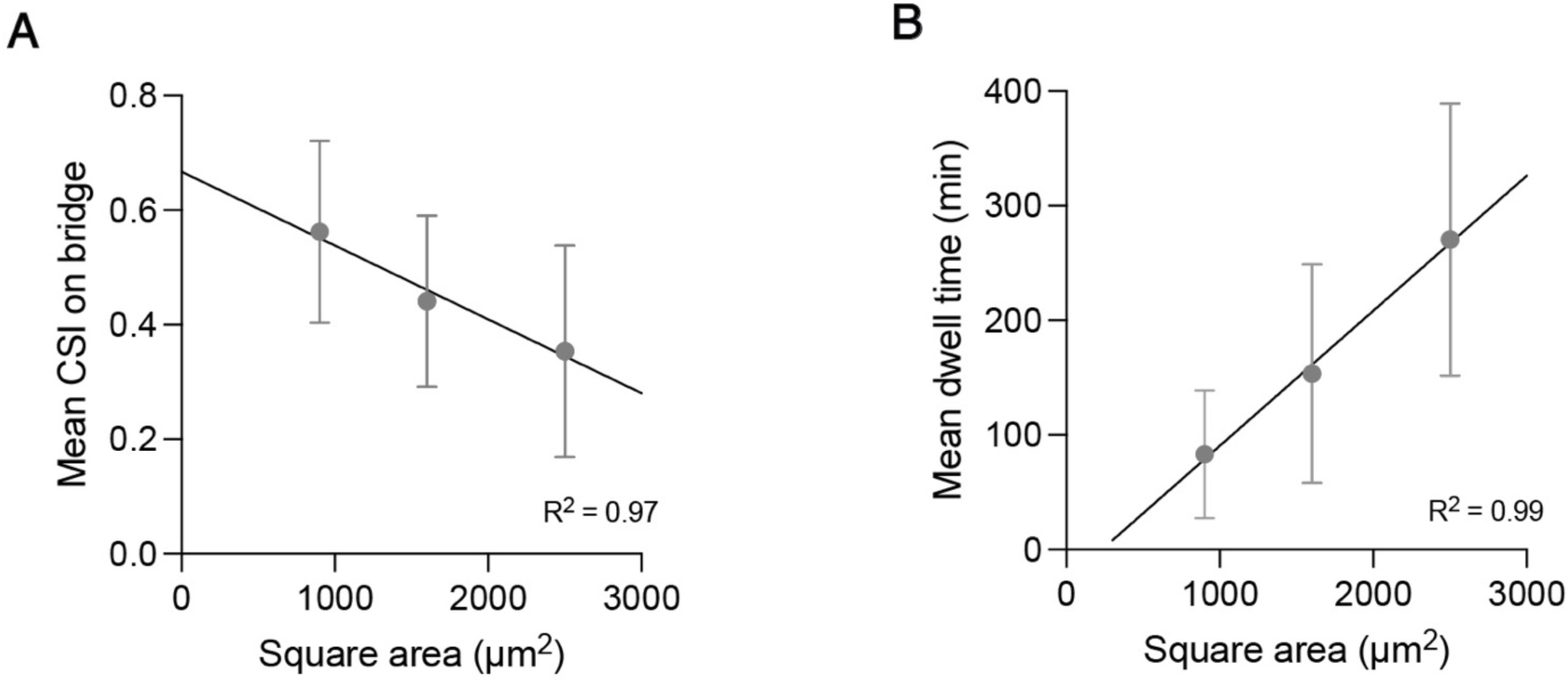
Relation between cell shape index, dwell time and square area. **(A)** Linear correlation between the mean cell shape index on bridge and the area of the square at each end. The regression coefficient is 0.9723. **(B)** Linear correlation between the mean dwell time on the square and the area of the square. The regression coefficient is 0.9951. Mean ± S.D.

### Supplementary Movies

**Supplementary Movie 1 –** Time-lapse sequences showing the back-and-forth motion of an individual MCF-10A epithelial cell on dumbbell-shaped fibronectin micropatterns with bridge lengths of 40 µm, 80 µm, 120 µm, and 160 µm and square ends measuring 40 µm x 40 µm. The cell is imaged in Differential Interference Contrast (DIC) mode, with the micropattern labeled with rhodamine-conjugated fibronectin and the nucleus stained with Hoechst. The scale bar represents 20 µm, and the frame rate is 3 minutes per frame.

**Supplementary Movie 2 –** Typical sequence of automatic tracking of an individual MCF-10A cell observed in DIC mode and analyzed using the FIJI plugin Cellpose 2.0, which provides the cell outline for each frame. This enables the determination of the cell’s spreading area, perimeter, and the coordinates of the cell front, rear, and center of mass.

**Supplementary Movie 3 –** Representative time-lapse sequences (t=20 hours) showing the elongated and compacted morphologies adopted by an individual MCF-10A epithelial cell during its back-and-forth motion on a dumbbell-shaped fibronectin micropattern with a bridge length of 160 µm. The cell is imaged using Differential Interference Contrast (DIC) microscopy, with the micropattern labeled with rhodamine-conjugated fibronectin and the nucleus stained with Hoechst. The scale bar represents 20 µm, and images were captured at 3-minute intervals.

**Supplementary Movie 4 –** Representative time-lapse sequences showing both a failed (t=24 hours) and a successful (t=3 hours and 51 minutes) crossing event during the back-and-forth motion of an individual MCF-10A epithelial cell on a dumbbell-shaped fibronectin micropattern with a bridge length of 160 µm. The cell is imaged using Differential Interference Contrast (DIC) microscopy, with the micropattern labeled with rhodamine-conjugated fibronectin and the nucleus stained with Hoechst. The scale bar represents 20 µm, and images were captured at 3-minute intervals.

**Supplementary Movie 5 –** Time-lapse sequences of actin retrograde flow in both elongated and compacted lamellipodia, observed in confocal mode using a 100× silicone objective (N.A. = 1.35). Images were acquired at a rate of one frame every 1.25 seconds for a total duration of five minutes.

**Supplementary Movie 6 –** High-resolution confocal Z-stack sequence (Nikon Spatial Array Confocal, NSPARC) of an elongated cell morphology, obtained using a galvano scanner at 60× magnification (Plan Apo oil immersion objective, N.A. 1.42) with a step size of 0.17 µm for three channels (DAPI, TRITC, and FITC). Actin filaments are shown in black, and DAPI is shown in blue.

**Supplementary Movie 7 –** High-resolution confocal Z-stack images (Nikon Spatial Array Confocal, NSPARC) of a compacted cell morphology, obtained using a galvano scanner at 60× magnification (Plan Apo oil immersion objective, N.A. 1.42) with a step size of 0.17 µm for three channels (DAPI, TRITC, and FITC). Actin filaments are shown in black, and DAPI is shown in blue.

**Supplementary Movie 8 –** Schematic movies of simulations of elongated (top), compacted (middle) and switching (bottom) cells moving through the confining dumbbell pattern. Color of the dot indicates state of polarization (orange: elongated, alpha=1; purple: compacted, alpha=-2), and arrow indicates the polarity of the cell. Plots on the right indicate simultaneous migration trajectories along the x-direction. The dumbbell-shaped micropattern is composed of a bridge of 160 µm in length and 6 µm in width connected at each extremity to a square of 40 µm × 40 µm.

**Supplementary Movie 9 –** Time-lapse sequences of a morphological switch from an elongated to a compacted state, captured in epifluorescence mode with actin filaments labeled using SPY555-FastAct™ (t = 24 hours) and in DIC mode (t = 24 hours). The epithelial cell (MCF10-A) migrates on a dumbbell-shaped fibronectin micropattern, featuring a bridge 160 µm in length and 6 µm in width, with square ends measuring 40 µm × 40 µm. The nucleus is stained blue with Hoechst. The scale bar represents 40 µm.

**Supplementary Movie 10 –** Time-lapse sequence in DIC mode of an epithelial cell (MCF10-A) migrating on fibronectin microstripe measuring 500 µm in length and 6 µm in width (t = 20 hours). The nucleus is stained blue with Hoechst. The scale bar represents 10 µm.

**Supplementary Movie 11 –** Successive time-lapse sequences in DIC mode of an individual epithelial cell (MCF10-A) migrating on a dumbbell-shaped fibronectin micropattern, featuring a bridge 160 µm in length and 6 µm in width, with square ends measuring 40 µm x 40 µm. Nocodazole-treated cells exhibit a loss of polarity, characterized by oscillations in the first sequence. In the second sequence, the cells fail to enter the bridge, and in the third sequence, they remain compacted while also failing to enter the bridge. In the first sequence, the cell is treated with 2.5 µM nocodazole at t = 1 hour and 30 minutes (t = 30 hours). In the second and third sequences, the cell is treated with 2.5 µM nocodazole at t = 2 hours and 30 minutes (t = 30 hours).

**Supplementary Movie 12 –** Time-lapse sequence in DIC mode of an epithelial cell (MCF10-A) migrating on a dumbbell-shaped fibronectin micropattern, featuring a bridge 160 µm in length and 6 µm in width, with square ends measuring 40 µm x 40 µm (t= 20 hours and 55 minutes). The cell is treated with 20 nM Latrunculin B at t= 03 hours and 50 minutes. The scale bar represents 10 µm.

**Supplementary Movie 13 –** Time-lapse sequence in DIC mode of an epithelial cell (MCF10-A) migrating on a dumbbell-shaped fibronectin micropattern, featuring a bridge 160 µm in length and 6 µm in width, with square ends measuring 40 µm x 40 µm (t= 21 hours and 40 minutes). The cell is treated with 10 µM Y27632 at t= 03 hours and 15 minutes. The scale bar represents 50 µm.

**Supplementary Movie 14 –** Time-lapse sequence in DIC mode of an epithelial cell (MCF10-A) migrating on a dumbbell-shaped fibronectin micropattern, featuring a bridge 160 µm in length and 6 µm in width, with square ends measuring 30 µm x 30 µm (t= 20 hours). The fibronectin micropattern is red and the nucleus in blue. The scale bar represents 20 µm.

**Supplementary Movie 15 –** Time-lapse sequence in DIC mode of an epithelial cell (MCF10-A) migrating on a dumbbell-shaped fibronectin micropattern, featuring a bridge 160 µm in length and 6 µm in width, with square ends measuring 50 µm x 50 µm (t = 20 hours). The fibronectin micropattern is depicted in red, and the nucleus is stained blue. The scale bar represents 20 µm.

**Supplementary Movie 16 –** Time-lapse sequence in DIC mode of an epithelial cell (MCF10-A) migrating on a dumbbell-shaped fibronectin micropattern, featuring a bridge 160 µm in length and 6 µm in width, with square ends measuring 50 µm x 50 µm (t = 20 hours). The cell is treated with 1 nM of Japlakinolide 30 min before the beginning of the timelapse experiment. The fibronectin micropattern is depicted in red, and the nucleus is stained blue. The scale bar represents 20 µm.

## Supplementary Theory Note

Our modeling approach has two key goals: first, to identify a minimal description of the polarity dynamics of elongated vs compacted cells that quantitatively captures the key experimental statistics. Second, to provide a framework to test the predicted statistics of various implementation of the dynamic switching between these to morphological states that we can compare to experiment. Together, these two aspects provide a quantitative description of the experimental data at the level of cell nucleus trajectories. Further theoretical work will be required to link the observed morphological switch to a more mechanistic description [1–3].

To achieve a minimal description of a confined migrating cell with a small number of parameters, we developed an active particle model that captures the essential features of the polarity dynamics and the interaction of the cell with the confining micropattern. Such active particle models for confined migration have been inferred from experiment [4] and derived from mechanistic models based on molecular clutch model assumption [3], making them a promising framework to describe the long time-scale dynamics of confined migrating cells. We first describe the implementation of elongated vs compacted states (section 1), followed by the implementation of the dynamical switch (section 3).

### 1 Model of migration dynamics of elongated and compacted cells

We focus on describing the motion of the cells in the *x*-direction along the long axis of the micropattern, as very little motion occurs in the orthogonal direction. The dimensions of the micropattern are given by the bridge length *L* and the island side length *a*, with the center of the bridge at *x* = 0. We consider the polarity of the cell to be described by a two-dimensional vector **p** = (*p_x_*, *p_y_*), since on the islands of the micropattern, the cell polarity may orient in 2D, allowing the cell to turn around. During the traversal of the bridge (−*L*/2 *< x < L*/2), we assume the cell polarity to be oriented along the *x*-axis, i.e. *p_y_* = 0.

The dynamics of cell position *x*(*t*) are then described by an overdamped equation of motion

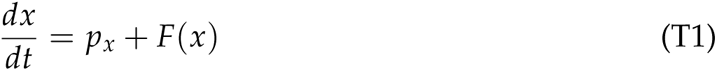

where *F*(*x*) are forces arising due to interaction with the micropattern boundary, for which we take a simple polynomial repulsive potential force:

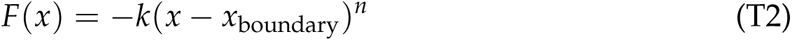

where we take *k* = 1 and *n* = 8 throughout, and *x*_boundary_ = ±(*a* + *L*/2).

The second component of our model are the description of the polarity dynamics of the cell, which we assume to be described by

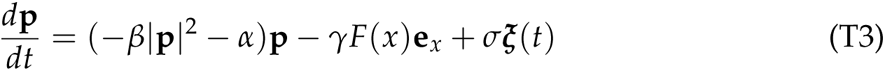

where **e***_x_* = (1, 0) is the unit vector along *x*, and ***ξ***(*t*) is a Gaussian white noise with ⟨***ξ***(*t*)⟩ = 0 and ⟨*ξ_µ_*(*t*)*ξ_ν_*(*t*^′^)⟩ = *δ_µν_δ*(*t* − *t*^′^).

The first term of Eq. (T3) is a standard formulation of active particle dynamics [5] including the two lowest order terms allowed by symmetry. Similar polarity dynamics have in previous work been inferred directly from experimental trajectories [4] and been derived from mechanistic considerations [3]. Taking *β >* 0 throughout, the sign of *α* controls the polarity dynamics of the cell.

For *α >* 0, the polarity dynamics are effectively a harmonic potential with a fixed point at **p** = 0. In this case, the polarity dynamics undergo a persistent random walk (with time-scale ~ 1/*α* for *β* → 0). Biologically, this corresponds to a state in which two protrusions left and right of the cell compete with each other, allowing rapid switches in the direction of motion of the cell center.

For α < 0, **p** = 0 becomes an unstable fixed point and a new stable fixed point at |**p**| = 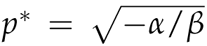 emerges. Biologically, this corresponds a single dominant protrusion, such that the overall cell polarity has a non-zero average and cannot instantaneously switch sign. Importantly, if the polarity dynamics are 2D, the angle of the polarity vector is a soft mode that can rotate freely, allowing reorientations of the cell. In our model, we assume this to be the case on the adhesive islands of the system. However, when the cell is highly confined into the 1D bridge connecting the islands, *p_y_* = 0, and therefore there is no soft mode in the system: for the cell to reorient, *p_x_* has to flip from +*p*^∗^ *p*^∗^ (or vice versa) by crossing the potential barrier at *p* = 0. This means that once confined to the channel, reorientations are rare, leading to highly persistent crossing of the bridge. Importantly, our assumption that the polarity dynamics are 2D on the islands also implies that highly persistent cells (*α <* 0) can remain polarized and rapidly reorient when they are on the island, due to the soft mode of the polarity dynamics.

Finally, we also include a term *γF*(*x*)**e***_x_* in our model, which causes the cell polarity to reorient when the cell reaches the boundaries of the micropattern. This is to avoid cells remaining highly polarized and moving against the micropattern boundary, which is not observed experimentally.

Based on initial parameter explorations, we take *β* = 10^−4^ and *σ* = 100 (both consistent with previously used parameters in an active particle model of confined cell migration [4]), and *γ* = 10. The qualitative behaviour of the system does not sensitively depend on the numerical value of these parameters, beyond their order of magnitude (see Table T1).

We then proceed to fit the value of *α*, on which the behaviour does depend sensitively.

We fit *α* separately for elongated (E) and compacted (C) states based on the experimentally measured average crossing speed of cells across the bridge, and find best fit values of *α_E_* = 1 and *α_C_*= − 2 (Fig. T1). This is in line with our initial hypothesis that elongated cells exhibit on average unpolarized behaviour corresponding to positive *α*, while compacted cells have negative values of *α*.

**Figure T1:**
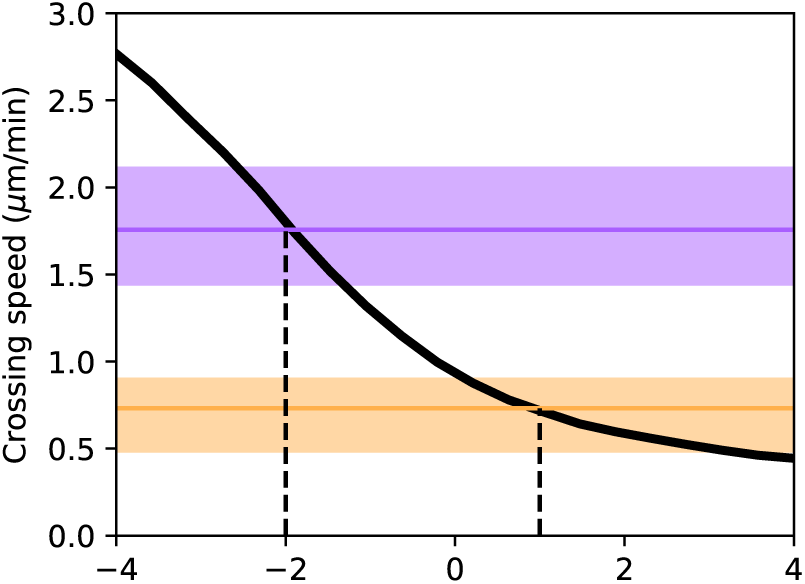
Bridge crossing speed as a function of *α*. Black line: simulations. Purple: measured value for compacted cells, orange: measured value for elongated cells. Shaded areas: interquartile range.

### 2 Inference of nonlinear polarity dynamics

To directly test the transition from negative to positive *α*, we use stochastic inference to infer the polarity dynamics of cells as they are transiting the center of the bridge. Based on our model, for *x* ≈ 0, we expect the dynamics to be described by

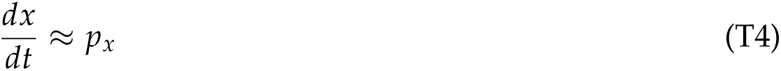

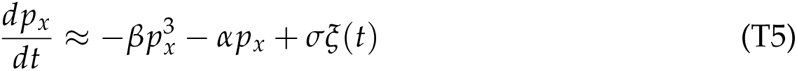

Differentiating the first equation, substituting the second, and introducing the cell velocity *v* = *dx*/*dt* as a variable, this is equivalent to underdamped dynamics of the form [6]

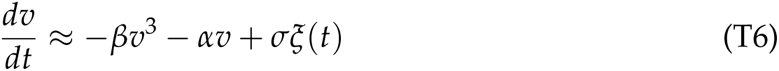

and therefore the conditional average of the acceleration should give direct access to the polarity dynamics, and thus *α* and *β* (see [7] for a note on caveats regarding effects of discretization):

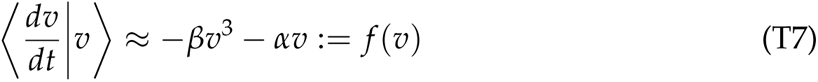

This is the quantity plotted in Fig. 2P of the main text, which compares well between model and experiment. Note that to deal with limited statistics, we make use of the known mirror symmetry *x x* of the micropattern, i.e. *f* (−*v*) = − *f* (*v*), and infer the symmetrized term *f* (*v*).

### 3 Model of the switching dynamics

Having established a minimal model for the dynamics of cells migrating in elongated and compacted states, we next explore the statistics of switches between these two states. Our aim here is to provide a minimal model to identify the statistics of the switching by comparison to experiment, rather than to provide a mechanistic model of the switches based on underlying biological mechanisms, which will be an important question to address in future work.

Based on the experimental observations, we simplify the morphological dynamics to have two discrete states (E, C), for which we explore different switching rules.

#### Uniform switching rates

We first explore a scenario where the stochastic switching between E and C states occurs at constant rates, independent of time and the location of the cell. For this, we use the Gillespie algorithm to generate the time-intervals spent in each state.

**Figure T2:**
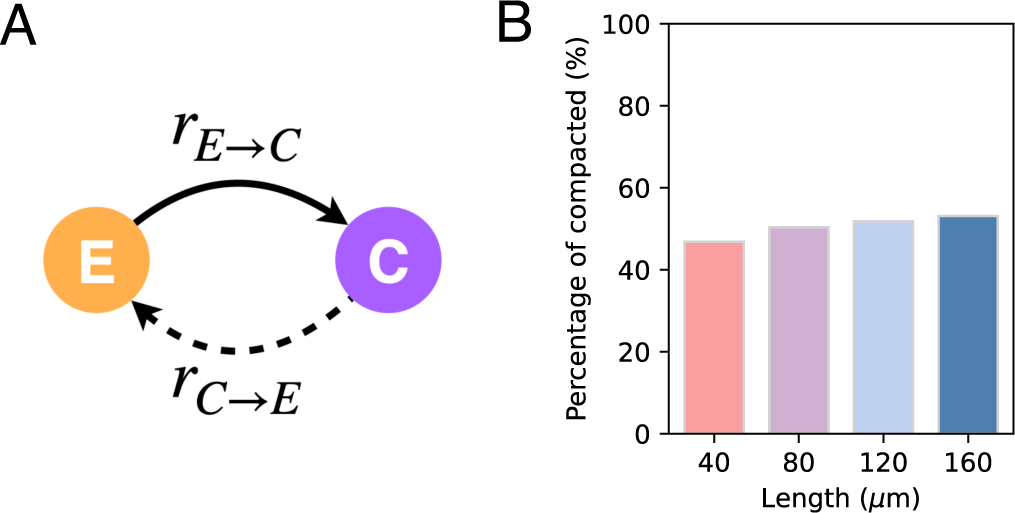
Prediction of the constant switching rate model. (A) Schematic of constant rate model. (B) Percentage of compacted cells on the bridge as a function of bridge length. Using *λ* = 1 h^−1^.

For a two-state system, the ratio of forward and backward rates *r_C_*_→*E*_ and *r_E_*_→*C*_, respectively (Fig. T2A), determines the occupation probabilities of each state: *P_E_*/*P_C_* = *r_C_*_→*E*_/*r_E_*_→*C*_. Since *P_E_* ≈ *P_C_* ≈ 0.5, *r_E_*_→*C*_ ≈ *r_C_*_→*E*_: = *λ*, where *λ* sets the overall time-scale of switching.

However, in this model, state switching is completely independent of the micropattern geometry. The fraction of compacted cells is therefore expected to be constant with bridge length (Fig. T2B), inconsistent with experimental observation (Fig. 3D).

#### Geometry-sensitive switching rates

Having ruled out the case of uniform switching rates, we hypothesized that the probability of switching between states is determined by the local confinement of the cell. A geometry adaptation of polarity dynamics has previously been inferred from experiments [4] and derived from mechanistic models based on molecular clutch model assumption [3]. However, in this case, the polarization parameter *α* was assumed to be a direct function of the local position, i.e. *α* = *α*(*x*). These previous work considered the limit of short bridges (*L <* 60*µ*m) and a different cell type.

Based on our observation that there are elongated cell states on the bridge, the data are not consistent with an instantaneous adaptation of the polarity dynamics. Furthermore, the observation of long-term memory of the morphological state across transitions suggests that cells can remain in a compacted morphology when they enter the island, which is consistent with our observation of lower spreading areas of cells on islands subsequent to a transition with a compacted morphology. This also suggests that geometry adaptation is not instantaneous.

Generalizing the arguments in [3, 4], we therefore first test the simplest implementation in which the switching rates are proportional to the local width of the micropattern. The local width *W*(*x*) of a dumbbell micropattern with bridge width *w*, bridge length *L* and square side length *a* is given by

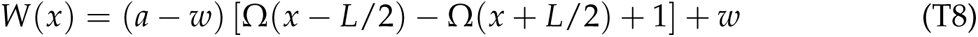

where Ω(*x*) = 1/(1 + *e*^−^*^x^*^/^*^ℓ^*) is a sigmoid function, and we use *ℓ* = 5*µ*m. We then take position-dependent rate functions *r_C_*_→*E*_(*x*) and *r_E_*_→*C*_(*x*),

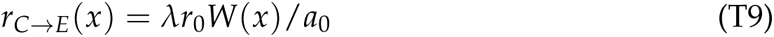

where we normalize the width by the standard square length *a*_0_ = 40*µ*m, such that the normalized width function *W*(*x*)/*a*_0_ is equal to 1 on the squares. The overall timescale of the switching process is then determined by *λ*. Furthermore, we take *f* (*x*) to have the opposite behaviour to *g*(*x*), i.e.

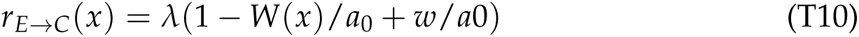

Note an extra factor of *r*_0_ is included that allows adjusting the relative proportions of elongated and compacted cells.

The overall time-scale of switching, *T* = 1/*λ*, determines how quickly cells adapt to the local environment. For *T* << *τ*, where *τ* is the dwell time spent on the island in between transitions, many switches occur during a single transition, and thus there is no memory of the local geometry across transitions. For *T* >> *τ*, the morphological state of the cell is strongly correlated from one transition to the next.

We demonstrate this by plotting the probability ratios *P*(*C C*)/*P*(*C*) and *P*(*E E*)/*P*(*E*) (Fig. T3). In the absence of memory across transitions, both fractions are equal to unity, since *P*(*C C*) = *P*(*C*) in that case, which we indeed observe for high *λ*, i.e. *T* << *τ*. Reducing *λ* below the transition rate 1/*τ* leads to an increase of the probability ratios, as *P*(*C*|*C*) *> P*(*C*), and similar for elongated states, indicating memory across transitions.

Note that the switching process in our model is Markovian (i.e. it only depends on the current state of the system). With ‘memory’, we refer to the time-scale of switches being larger than the time-scale of transitions, indicating that the cell can maintain its morphological state despite changing local geometry, i.e. maintain a memory of its state.

**Figure T3:**
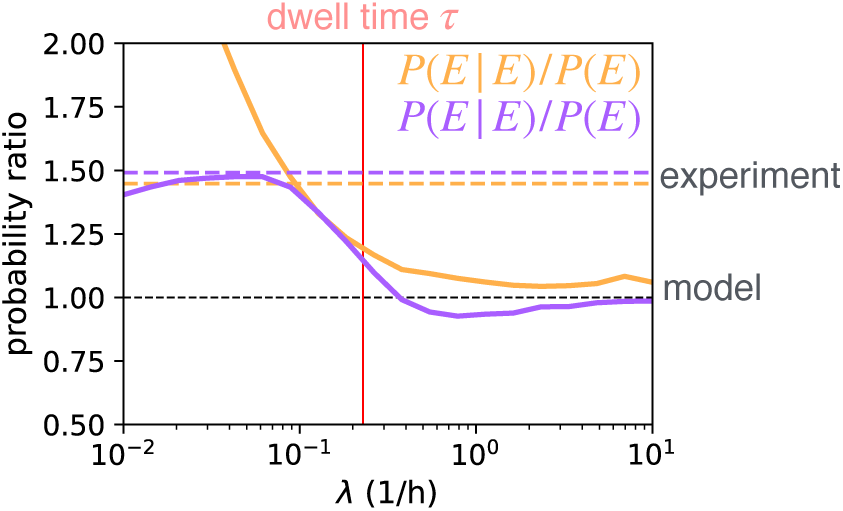
Probability ratios indicating the presence of memory across transitions as a function of the average switching rate. Horizontal dashed lines: probability ratios observed experimentally. Solid lines: predicted probability ratios as a function of *λ*. Vertical red line: experimentally measured average dwell time.

### 4 Numerical implementation and parameter overview

We numerically integrate the stochastic dynamical system governing the migration, polarity and state dynamics using custom-written python code using numpy [8]. We use a stochastic Euler scheme with small time-step *δt*:

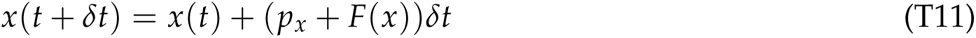

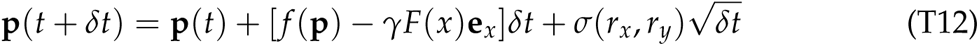

where *r_x_*_,*y*_ ∼ *N*(0, 1) are zero-mean, unit-variance normally distributed random numbers. We use a time interval *δt* = 0.005h, iterated for *N_t_* = 4000 time steps, equating to a total trajectory length of 20h like in the experiment. The parameters used throughout the figures are specified in Table T1.

**Table T1:**
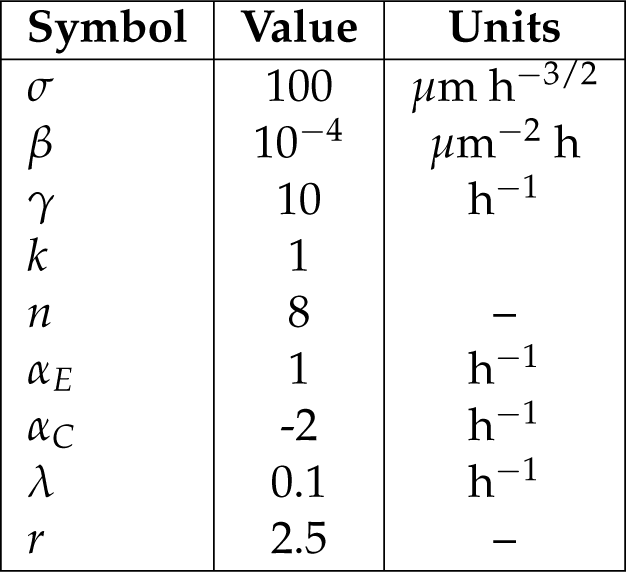
Overview of parameters used.

## Notes

### Competing Interest Statement

The authors have declared no competing interest.

