## Supplementary Theory Note for "The actin cortex acts as a mechanical memory of morphology in confined migrating cells"

The dynamics of cell position $x(t)$ are then described by an overdamped equation of motion

$$\frac{dx}{dt} = p_x + F(x) \tag{T1}$$

where $F(x)$ are forces arising due to interaction with the micropattern boundary, for which we take a simple polynomial repulsive potential force:

$$F(x) = -k(x - x_{\text{boundary}})^n \tag{T2}$$

where we take $k = 1$ and $n = 8$ throughout, and $x_{\text{boundary}} = \pm(a + L/2)$.

The second component of our model are the description of the polarity dynamics of the cell, which we assume to be described by

$$\frac{d\mathbf{p}}{dt} = (-\beta|\mathbf{p}|^2 - \alpha)\mathbf{p} - \gamma F(x)\mathbf{e}_x + \sigma\boldsymbol{\xi}(t) \tag{T3}$$

where $\mathbf{e}_x = (1, 0)$ is the unit vector along $x$, and $\boldsymbol{\xi}(t)$ is a Gaussian white noise with $\langle\boldsymbol{\xi}(t)\rangle = 0$ and $\langle\xi_\mu(t)\xi_\nu(t')\rangle = \delta_{\mu\nu}\delta(t - t')$.

The first term of Eq. (T3) is a standard formulation of active particle dynamics [5] including the two lowest order terms allowed by symmetry. Similar polarity dynamics have in previous work been inferred directly from experimental trajectories [4] and been derived from mechanistic considerations [3]. Taking $\beta > 0$ throughout, the sign of $\alpha$ controls the polarity dynamics of the cell.

For $\alpha > 0$, the polarity dynamics are effectively a harmonic potential with a fixed point at $\mathbf{p} = 0$. In this case, the polarity dynamics undergo a persistent random walk (with time-scale $\sim 1/\alpha$ for $\beta \to 0$). Biologically, this corresponds to a state in which two protrusions left and right of the cell compete with each other, allowing rapid switches in the direction of motion of the cell center.

For $\alpha < 0$, $\mathbf{p} = 0$ becomes an unstable fixed point and a new stable fixed point at $|\mathbf{p}| = p^* = \sqrt{-\alpha/\beta}$ emerges. Biologically, this corresponds a single dominant protrusion, such that the overall cell polarity has a non-zero average and cannot instantaneously switch sign. Importantly, if the polarity dynamics are 2D, the angle of the polarity vector is a soft mode that can rotate freely, allowing reorientations of the cell. In our model, we assume this to be the case on the adhesive islands of the system. However, when the cell is highly confined into the 1D bridge connecting the islands, $p_y = 0$, and therefore there is no soft mode in the system: for the cell to reorient, $p_x$ has to flip from $+p^* \to -p^*$ (or vice versa) by crossing the potential barrier at $p = 0$. This means that once confined to the channel, reorientations are rare, leading to highly persistent crossing of the bridge. Importantly, our assumption that the polarity dynamics are 2D on the islands also implies that highly persistent cells ($\alpha < 0$) can remain polarized and rapidly reorient when they are on the island, due to the soft mode of the polarity dynamics.

Finally, we also include a term $\gamma F(x)\mathbf{e}_x$ in our model, which causes the cell polarity to reorient when the cell reaches the boundaries of the micropattern. This is to avoid cells remaining highly polarized and moving against the micropattern boundary, which is not observed experimentally.

Based on initial parameter explorations, we take $\beta = 10^{-4}$ and $\sigma = 100$ (both consistent with previously used parameters in an active particle model of confined cell migration [4]), and $\gamma = 10$. The qualitative behaviour of the system does not sensitively depend on the numerical value of these parameters, beyond their order of magnitude (see Table T1).

We then proceed to fit the value of $\alpha$, on which the behaviour does depend sensitively.

We fit $\alpha$ separately for elongated (E) and compacted (C) states based on the experimentally measured average crossing speed of cells across the bridge, and find best fit values of $\alpha_E = 1$ and $\alpha_C = -2$ (Fig. T1). This is in line with our initial hypothesis that elongated cells exhibit on average unpolarized behaviour corresponding to positive $\alpha$, while compacted cells have negative values of $\alpha$.

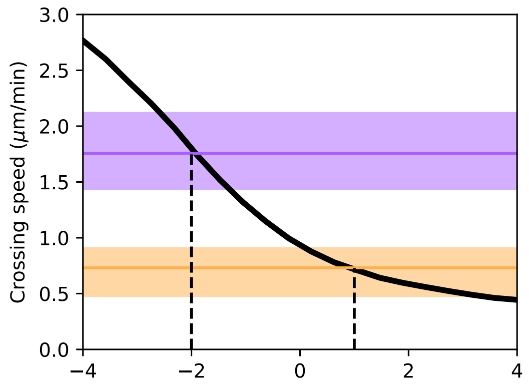


Figure T1: **Bridge crossing speed as a function of $\alpha$.** Black line: simulations. Purple: measured value for compacted cells, orange: measured value for elongated cells. Shaded areas: interquartile range.

### 2 Inference of nonlinear polarity dynamics

To directly test the transition from negative to positive $\alpha$, we use stochastic inference to infer the polarity dynamics of cells as they are transiting the center of the bridge. Based on our model, for $x \approx 0$, we expect the dynamics to be described by

$$\frac{dx}{dt} \approx p_x \tag{T4}$$

$$\frac{dp_x}{dt} \approx -\beta p_x^3 - \alpha p_x + \sigma \xi(t) \tag{T5}$$

Differentiating the first equation, substituting the second, and introducing the cell velocity $v = dx/dt$ as a variable, this is equivalent to underdamped dynamics of the form [6]

$$\frac{dv}{dt} \approx -\beta v^3 - \alpha v + \sigma \xi(t) \tag{T6}$$

and therefore the conditional average of the acceleration should give direct access to the polarity dynamics, and thus $\alpha$ and $\beta$ (see [7] for a note on caveats regarding effects of discretization):

$$\left\langle \frac{dv}{dt} \middle| v \right\rangle \approx -\beta v^3 - \alpha v := f(v) \tag{T7}$$

This is the quantity plotted in Fig. 2P of the main text, which compares well between model and experiment. Note that to deal with limited statistics, we make use of the known mirror symmetry $x \to -x$ of the micropattern, i.e. $f(-v) = -f(v)$, and infer the symmetrized term $f(v)$.

### 3 Model of the switching dynamics

Having established a minimal model for the dynamics of cells migrating in elongated and compacted states, we next explore the statistics of switches between these two states. Our aim here is to provide a minimal model to identify the statistics of the switching by comparison to experiment, rather than to provide a mechanistic model of the switches based on underlying biological mechanisms, which will be an important question to address in future work.

Based on the experimental observations, we simplify the morphological dynamics to have two discrete states (E, C), for which we explore different switching rules.

**Uniform switching rates**: We first explore a scenario where the stochastic switching between E and C states occurs at constant rates, independent of time and the location of the cell. For this, we use the Gillespie algorithm to generate the time-intervals spent in each state.

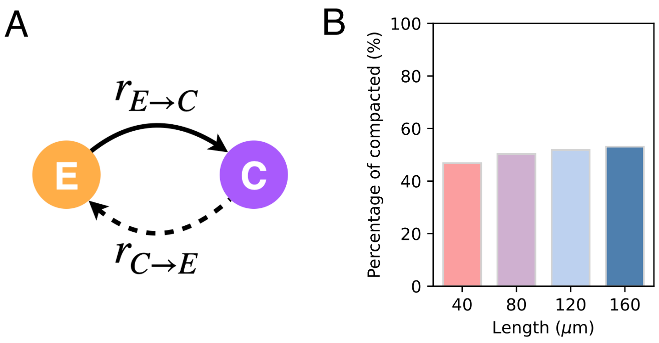


Figure T2: **Prediction of the constant switching rate model.** (A) Schematic of constant rate model. (B) Percentage of compacted cells on the bridge as a function of bridge length. Using $\lambda = 1\,\mathrm{h}^{-1}$.

For a two-state system, the ratio of forward and backward rates $r_{C\to E}$ and $r_{E\to C}$, respectively (Fig. T2A), determines the occupation probabilities of each state: $P_E/P_C = r_{C\to E}/r_{E\to C}$. Since $P_E \approx P_C \approx 0.5$, $r_{E\to C} \approx r_{C\to E} := \lambda$, where $\lambda$ sets the overall timescale of switching.

However, in this model, state switching is completely independent of the micropattern geometry. The fraction of compacted cells is therefore expected to be constant with bridge length (Fig. T2B), inconsistent with experimental observation (Fig. 3D).

**Geometry-sensitive switching rates**

Having ruled out the case of uniform switching rates, we hypothesized that the probability of switching between states is determined by the local confinement of the cell. A geometry adaptation of polarity dynamics has previously been inferred from experiments [4] and derived from mechanistic models based on molecular clutch model assumption [3]. However, in this case, the polarization parameter $\alpha$ was assumed to be a direct function of the local position, i.e. $\alpha = \alpha(x)$. These previous work considered the limit of short bridges ($L < 60\mu$m) and a different cell type.

Based on our observation that there are elongated cell states on the bridge, the data are not consistent with an instantaneous adaptation of the polarity dynamics. Furthermore, the observation of long-term memory of the morphological state across transitions suggests that cells can remain in a compacted morphology when they enter the island, which is consistent with our observation of lower spreading areas of cells on islands subsequent to a transition with a compacted morphology. This also suggests that geometry adaptation is not instantaneous.

Generalizing the arguments in [3,4], we therefore first test the simplest implementation in which the switching rates are proportional to the local width of the micropattern. The local width $W(x)$ of a dumbbell micropattern with bridge width $w$, bridge length $L$ and square side length $a$ is given by

$$W(x) = (a - w)\left[\Omega(x - L/2) - \Omega(x + L/2) + 1\right] + w \tag{T8}$$

where $\Omega(x) = 1/(1 + e^{-x/\ell})$ is a sigmoid function, and we use $\ell = 5\mu$m.

We then take position-dependent rate functions $r_{C\to E}(x)$ and $r_{E\to C}(x)$,

$$r_{C\to E}(x) = \lambda r_0 W(x)/a_0 \tag{T9}$$

where we normalize the width by the standard square length $a_0 = 40\mu$m, such that the normalized width function $W(x)/a_0$ is equal to 1 on the squares. The overall timescale of the switching process is then determined by $\lambda$. Furthermore, we take $f(x)$ to have the opposite behaviour to $g(x)$, i.e.

$$r_{E\to C}(x) = \lambda(1 - W(x)/a_0 + w/a0) \tag{T10}$$

Note an extra factor of $r_0$ is included that allows adjusting the relative proportions of elongated and compacted cells.

The overall time-scale of switching, $T = 1/\lambda$, determines how quickly cells adapt to the local environment. For $T \ll \tau$, where $\tau$ is the dwell time spent on the island in between transitions, many switches occur during a single transition, and thus there is no memory of the local geometry across transitions. For $T \gg \tau$, the morphological state of the cell is strongly correlated from one transition to the next.

We demonstrate this by plotting the probability ratios $P(C|C)/P(C)$ and $P(E|E)/P(E)$ (Fig. T3). In the absence of memory across transitions, both fractions are equal to unity, since $P(C|C) = P(C)$ in that case, which we indeed observe for high $\lambda$, i.e. $T \ll \tau$. Reducing $\lambda$ below the transition rate $1/\tau$ leads to an increase of the probability ratios, as $P(C|C) > P(C)$, and similar for elongated states, indicating memory across transitions.

Note that the switching process in our model is Markovian (i.e. it only depends on the current state of the system). With 'memory', we refer to the time-scale of switches being larger than the time-scale of transitions, indicating that the cell can maintain its morphological state despite changing local geometry, i.e. maintain a memory of its state.

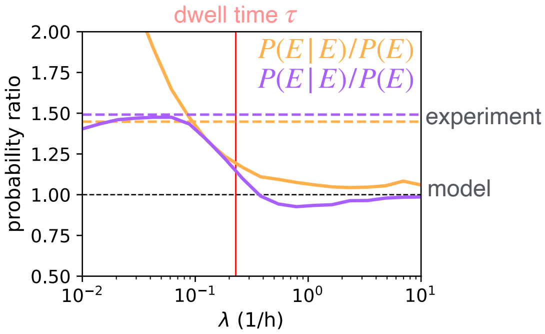


Figure T3: **Probability ratios indicating the presence of memory across transitions as a function of the average switching rate.** Horizontal dashed lines: probability ratios observed experimentally. Solid lines: predicted probability ratios as a function of $\lambda$. Vertical red line: experimentally measured average dwell time.

### 4 Numerical implementation and parameter overview

We numerically integrate the stochastic dynamical system governing the migration, polarity and state dynamics using custom-written python code using numpy [8]. We use a stochastic Euler scheme with small time-step $\delta t$:

$$x(t + \delta t) = x(t) + (p_x + F(x))\delta t \tag{T11}$$

$$\mathbf{p}(t + \delta t) = \mathbf{p}(t) + [f(\mathbf{p}) - \gamma F(x)\mathbf{e}_x]\delta t + \sigma(r_x, r_y)\sqrt{\delta t} \tag{T12}$$

where $r_{x,y} \sim \mathcal{N}(0, 1)$ are zero-mean, unit-variance normally distributed random numbers. We use a time interval $\delta t = 0.005$h, iterated for $N_t = 4000$ time steps, equating to a total trajectory length of 20h like in the experiment. The parameters used throughout the figures are specified in Table T1.

| Symbol | Value | Units |
| --- | --- | --- |
| $\sigma$ | 100 | $\mu$m h$^{-3/2}$ |
| $\beta$ | $10^{-4}$ | $\mu$m$^{-2}$ h |
| $\gamma$ | 10 | h$^{-1}$ |
| $k$ | 1 | |
| $n$ | 8 | – |
| $\alpha_E$ | 1 | h$^{-1}$ |
| $\alpha_C$ | -2 | h$^{-1}$ |
| $\lambda$ | 0.1 | h$^{-1}$ |
| $r$ | 2.5 | – |

Table T1: Overview of parameters used.
